# Discrete-to-analog signal conversion in human pluripotent stem cells

**DOI:** 10.1101/2021.11.05.467377

**Authors:** Laura Prochazka, Yale S. Michaels, Charles Lau, Mona Siu, Ting Yin, Diana Wu, Esther Jang, Ross D. Jones, Mercedes Vázquez-Cantú, Penney M. Gilbert, Himanshu Kaul, Yaakov Benenson, Peter W. Zandstra

**Author notes:** Corresponding Author: Peter W. Zandstra, Professor and Director, School of Biomedical Engineering, Director, Michael Smith Laboratories, University of British Columbia, Vancouver Campus, 2185 East Mall, Vancouver, BC Canada V6T 1Z4.

## Abstract

During development, state transitions are coordinated through changes in the identity of molecular regulators in a cell state- and dose specific manner. The ability to rationally engineer such functions in human pluripotent stem cells (hPSC) will enable numerous applications in regenerative medicine. Herein we report the generation of synthetic gene circuits that can detect a discrete cell state, and upon state detection, produce fine-tuned effector proteins in a programmable manner. Effectively, these gene circuits convert a discrete (digital-like) cell state into an analog signal by merging AND-like logic integration of endogenous miRNAs (classifiers) with a miRNA-mediated output fine-tuning technology (miSFITs). Using an automated miRNA identification and model-guided circuit optimization approach, we were able to produce robust cell state specific and graded output production in undifferentiated hPSC. We further finely controlled the levels of endogenous BMP4 secretion, which allowed us to document the effect of endogenous factor secretion in comparison to exogenous factor addition on early tissue development using the hPSC-derived gastruloid system. Our work provides the first demonstration of a discrete-to-analog signal conversion circuit operating in living hPSC, and a platform for customized cell state-specific control of desired physiological factors, laying the foundation for programming cell compositions in hPSC-derived tissues and beyond.

**Graphical Abstract:** 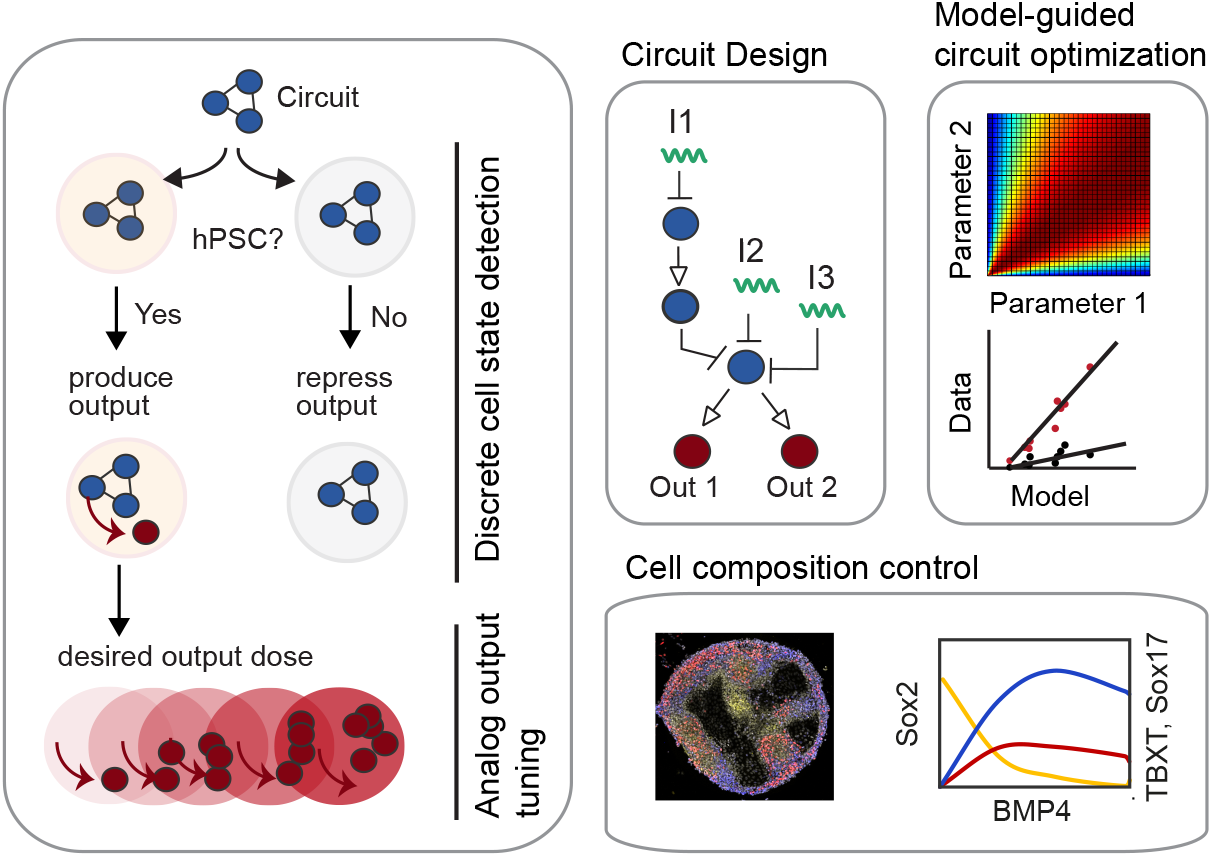

## Introduction

Robust gene-regulatory programs enable stem cells to self-renew and differentiate by sensing and responding to stimuli in a defined manner [1–3]. Crucially, these regulatory circuits are capable of integrating multiple internal and external input signals to achieve a high degree of specificity, resulting in lineage or cell state-specific activation of effector molecules [4, 5]. The production of effector molecules is often graded, or analog, where defined doses can lead to desirable proportions of downstream lineages [6–8]. The ability to engineer such gene-regulatory circuits into human pluripotent stem cells (hPSC) *de novo* would enable efficient production of desired cell types or tissues for research and regenerative medicine applications [9–13]. With the goal of controlling human cell function, substantial effort has been directed over the past two decades towards synthetic gene circuit engineering [14–21], with recent exciting developments using hPSC [22–26].

The majority of gene circuits implemented in human cells are logic gene circuits [27–32]. A handful of them have been designed to detect cell-internal endogenous input signals, enabling restriction of circuit action to desired cell types or cell states [33–38]. Here we define a cell state discrete, if it can be clearly discriminated from other cell states on the basis of a predefined set of molecular inputs that are detected and integrated by a circuit. The underlying circuit integrates the inputs in a function that can approximated by Boolean logic and autonomously “decides” if a desired downstream molecule, the output, is produced (On) or not (Off). One type of circuit that allows such discrete cell state discrimination are the so-called cell “classifiers” [35, 39]. Cell classifiers have been designed to detect and logically integrate endogenous microRNAs (miRNAs) and have proven useful for a variety of applications such as the specific killing of cancer cells [35, 40] or for screening miRNA drug candidates [41]. Additionally, single endogenous miRNAs have been employed to regulate synthetic genes to discriminate hPSCs from differentiated cells [42], for selection of PSC-derived mature cell types [36], or reprogrammed hPSC [43] and for specific killing of hPSC [36, 44]. Interestingly, endogenous miRNAs have also been exploited to fine tune expression levels of synthetic and natural genes in human cells [45]. Such graded or analog production of proteins is crucial for many applications where precise intervention of physiological behavior is required [45].

Despite this progress, current implementations of cell classifiers result in arbitrary On and Off levels that are highly dependent on parameters such as the promoter strength and delivery system, and thus are difficult to tune to the desired dose [34, 35, 46, 47]. Furthermore, miRNA-based systems implemented in stem cells typically operate with a single miRNA input and a single protein output [36, 42–44], limiting their applications. To date, no circuit has been reported that allows precise tuning of multiple desired proteins from desired discrete cell states, a function that stem cells perform continuously during development and would enable powerful control over stem cell differentiation.

Here we describe the design- and implementation strategy of synthetic gene circuits that are capable of performing cell state specific control of desired proteins in hPSC. Mechanistically, our system integrates an input module that uses discrete logic gate computations [35] to detect a set of hPSC-specific miRNAs, with an output module that operates with miRNA silencing-mediated fine tuners (miSFITs) that allows to pre-select for desired output dose [45]. Using a model-guided combinatorial screening approach, we first optimize circuit performance in hPSCs and then validate accurate discrete-to-analog signal processing for up to two fluorescent reporters. We further demonstrate the utility of this system for the autonomous induction of BMP4 dose-mediated hPSC microtissue patterning to achieve control over the proportions of differentiated cell types. This technology lays the foundation for rapid, model-guided engineering of cell-state specific analog circuits in hPSC, hPSC-derived cell types, and other human cells.

### Circuit design

To create discrete-to-analog converters, we designed a generic circuit that first detects a pluripotent state, restricting circuit actuation to undifferentiated hPSC and then tunes multiple protein outputs to defined doses (Figure 1A). Our design uses a bow-tie architecture [34] that allows decomposition of the circuit into two modules: a logic multi-input module, allowing discrete cell state detection by recognizing a computationally identified set of miRNAs; and an analog multi-output module, that tunes output dose to desired levels by pre-selection from a library of miRNA mediated fine-tuners (miSFITs) [45] (Figure 1B).

**Figure 1.**
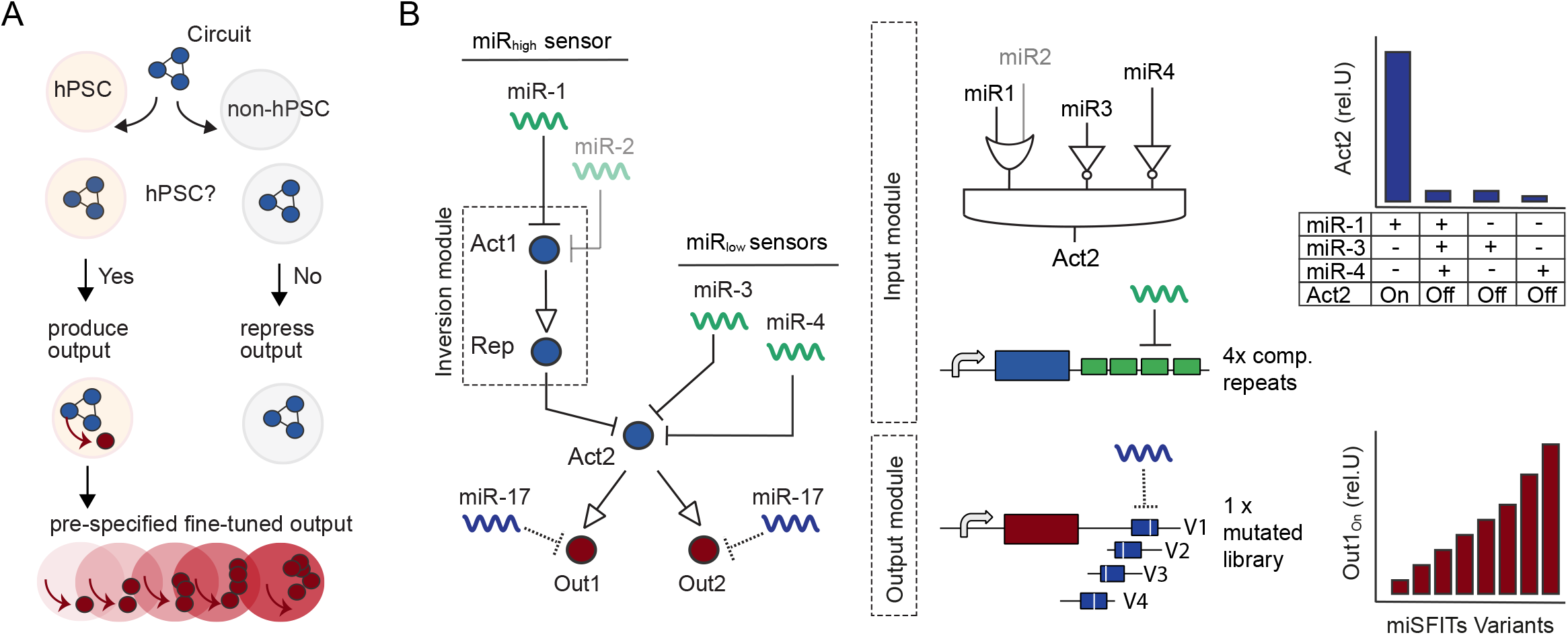
Circuit design. **A)** Schematic of circuit performance on cellular level. **B)** Generic circuit architecture of the discrete-to-analog converter (left). Logic input module (top) and analog output module (bottom). Act1 and Act2 are two different orthogonal synthetic transactivators, Rep is a synthetic miRNA-based repressor. Depicted are one miRhigh sensor (inversion module) that recognizes one or two miRNAs (miR1, miR2 = OR gate) and two miRlow sensors each recognizing one miRNA (miR3, miR4). To achieve discrete cell state actuation, sensors are built with four fully complementary repeats for miRNA recognition (targeting either Act1 in miRhigh sensors or Act2 in miRlow sensors) leading to Output production only if input combinations match (bar chart, right top). The performance for the depicted input module can be approximated with a Boolean logic function Output = miR1 OR miR2 AND NOT miR3 AND NOT miR4. The output module is controlled by Act2 and shows two protein outputs that are further controlled by miSFITs, a miR-17-based target library. The library uses 1x target sites containing various mutations to achieve fine-tuned levels of the output by variant selection (bar chart, right bottom).

The input module is composed of two kinds of miRNA sensors, here named miR_high_ - and miR_low_ sensors (Figure 1B). miR_low_ sensors directly repress a synthetic transactivator, Act2, by placing four fully complementary repeats of a given miRNA in the 3‘ UTR of Act2, resulting in Act2 expression only when the given miRNA is absent or at low levels. miR_high_ sensors are double inversion modules, where the endogenous miRNA represses an Act1-inducible repressor (here a synthetic miRNA) [35, 46] and thus results in high Act2 expression only if the given miRNA is highly expressed. A miR_high_ sensor can also be targeted by two or more miRNAs, forming an OR gate, that can improve the inversion performance and make it more robust to fluctuations in endogenous miRNAs [23, 39] (Figure 1B). Because all sensors converge on Act2, the integration of the miRNA signals can be approximated in an AND-like logic function [34]. This means that Act2 is produced at high levels only if highly expressed miRNAs are recognized by the miR_high_ sensors AND when miRNAs that are not expressed or active in a given cell state are recognized by miR_low_ sensors. If one of the miRNA inputs substantially differs to this profile, or multiple miRNA differ slightly, Act2 expression should be significantly repressed [34, 35], thereby enabling discrete cell state detection.

The analog output module is controlled by Act2 and thus, indirectly, by the endogenous miRNAs. Act2 can activate one or multiple protein outputs (Figure 1B). To fine-tune the expression levels of each output, we applied miSFITs, a previously reported approach that operates on a library of mutated human miRNA-17 target sites, placed as a single repeat in the 3’UTR of a desired gene (here protein outputs) (Figure 1B). Because miR-17 is ubiquitously and strongly expressed among most human cell types, including hPSC [48], mutations in the target regions lead to reduced binding strength of endogenous miR-17. The repression strength varies depending on the position and identity of the mismatched nucleotides [45]. Because of the high number of different mutant variants that can be created, expression dose can be tuned to very fine levels (Figure 1B, lower right panel). Thus, by selection of a variant from a library, circuits can be created that can produce a protein at any desired pre-specified or analog level [45].

## Results

### Automated identification of hPSC-specific miRNAs

In order to restrict circuit action to discrete cell states or cell types, a set of endogenous miRNAs must be identified to allow clear discrimination of the cell-state of interest (positive samples, here hPSC) from the other cell states (negative samples, here hPSC-derived differentiated cells) within a testing space. To address the challenge of manually selecting such a set of miRNAs, we have applied and further modified a previously developed computational platform that automates the miRNA candidate search and circuit design procedure [39]. This platform uses a mechanistic mathematical model that predicts circuit output production from miRNA expression data by seeking a set of miRNAs and underlying circuit with the largest classification margin (cMargin), or in other words, the largest fold change in calculated circuit output levels between positive samples and negative samples [39].

In order to apply the platform to identify hPSC-specific miRNAs from different published miRNA sequencing sources, we have modified the miRNA pre-selection step of the algorithm to allow selection of miRNA sequences instead of miRNA names (see Methods). Using three different data sets, covering four hPSC lines and 15 hPSC-derived cell states (Figure 2A, Table S1) [48–50], we identified a minimal set of miRNAs that allowed discrimination of hPSC from the other cell states with a cMargin of 1.16 corresponding to an average of 14.4-fold change between the hPSC group and the differentiated group (Figure 2A, Table S2). The algorithm identified three miRNAs: miRNA-302a, which is highly expressed in hPSC and plays a critical role in maintenance of the pluripotent state [49]; and miR-489 and miR-375, that are not expressed in hPSC but are expressed at different levels in the negative samples (Figure 2A, Table S2). miR-375 is a key regulator during differentiation of hPSC towards pancreatic progenitors and mature beta- and alpha-cells [48]. The role of miR-489 has been described in cancer but not, to the best of our knowledge, in hPSCs or during development. The underlying logic function the circuit performs can be approximated with: Output = miR-302a AND NOT miR-489 AND NOT miR-375 (Figure 2A, right). By increasing the maximal circuit input number constraints and / or considering the unpruned circuit version, we found additional circuits with slightly improved performance (Figure S1A). All circuits include miR-302a, miR-375 and miR-489 amongst other miRNAs (Figure S1A), supporting the importance of the three miRNAs for hPSC classification.

**Figure 2.**
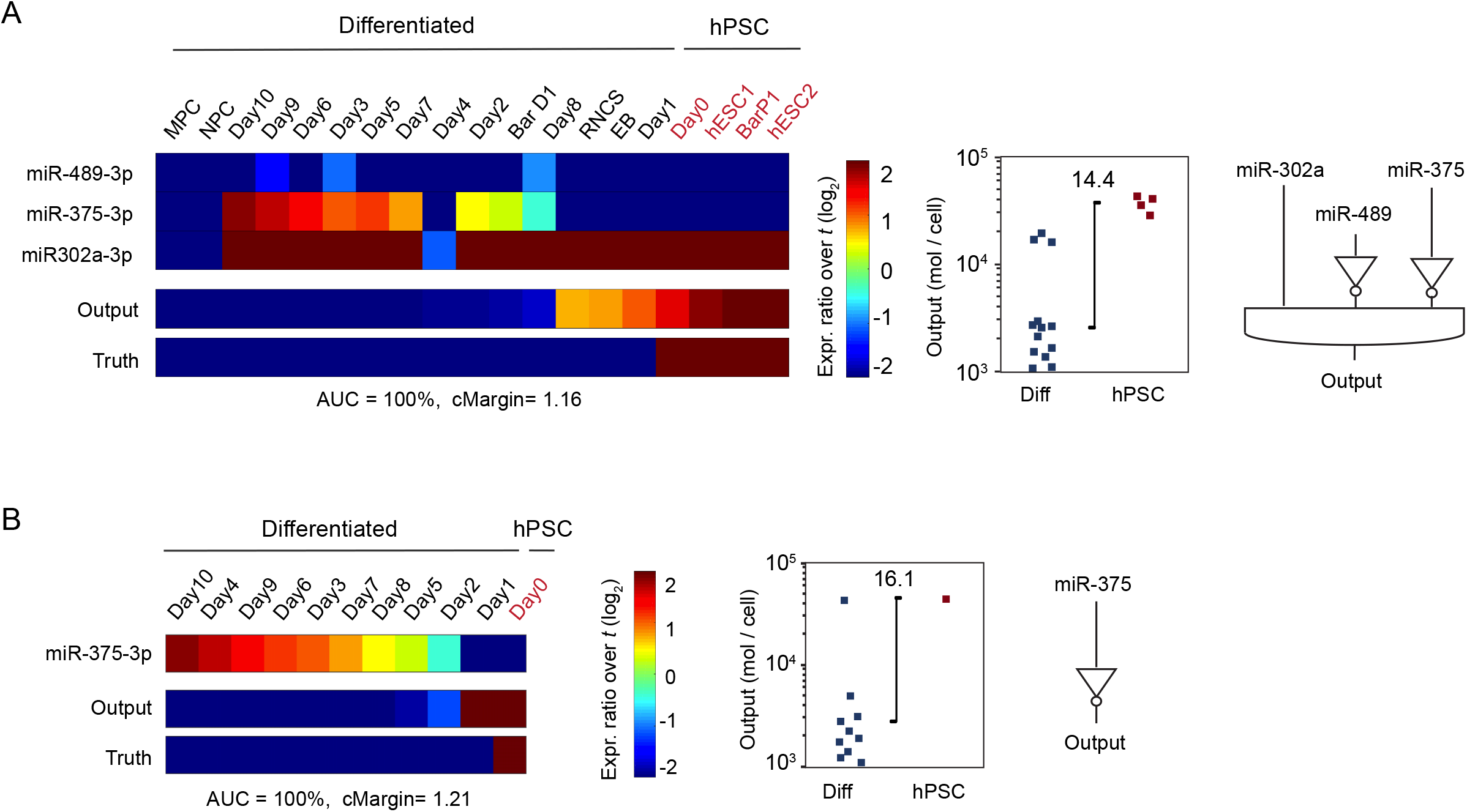
Automated identification of hPSC-specific miRNAs. **A)** Summary of the computationally identified pruned circuit showing input miRNAs and predicted circuit output levels using miRNA data sets from Fogel et al., Lipchina et al and Bar et al **(A)** or only Fogel et al data **(B)**. For both A and B) a constraint of maximum three inputs have been used. Expression levels of identified miRNAs inputs are given as fold change over the pre-set input abundance threshold (t) of the total miRNA pool (where t = 1%) (left). Calculated circuit output levels are given as mol / cell (middle) and logic connectivity of the identified miRNA is depicted (right). miRNA expression data and nomenclature in Table S1, raw output data and constraint files of the algorithm in Table S2 for A and Table S3 for B. See also Figure S1.

We further analyzed how well hPSC could be discriminated from committed cells using only the Fogel et al. [48] data set, which collected miRNA expression data from each day during differentiation towards pancreatic progenitors (Figure 2B). We found that miR-375 alone allowed discrimination of hPSC from the remaining differentiated cells (particularly by day 10 of differentiation). Cells measured one day after differentiation (day 1) showed only slightly reduced output production compared to hPSC and discrimination could be slightly improved with the addition of miR-489 and miR-708 (Figure S1B). In contrast, cells appearing as early as two days after differentiation (day 2), which adopt a definitive endoderm lineage, could be clearly discriminated from hPSC with miR-375 alone (Figure 2B, Table S3).

To sum up, we identified, in an automated manner, a three-input miRNA profile capable of discriminating hPSC from various differentiated cells, and a single miRNA (miR-375) to discriminate hPSC from cells differentiating to pancreatic progenitors. More broadly, our miRNA selection algorithm can be applied to the optimal selection of logic gates for cell state discrimination during hPSC differentiation.

### Endogenous miRNAs allow discrete and analog control of synthetic genes in hPSC

The expression level of a given miRNA is an important determinant of its ability to repress a synthetic gene. The repression strength is also dependent on the design of the target sites with respect to the number of repetitive miRNA target units and its surrounding sequence. Therefore, before implementing our discrete-to-analog converter, we investigated if computationally identified miRNAs (miR-302a, miR-375 and miR-489) and the fine tuner miRNA (miR-17) showed expected activity in hPSC. After confirming that our transfection procedure and / or the miRNA sensors do not substantially affect the pluripotency state, endoderm differentiation potential and viability of hPSC (Figure S2A-D), we analyzed the repression strength of each identified miRNA using a previously reported bi-directional miRNA reporter construct [35] (Figure 3A). Upon transient transfection of the sensor system into hPSC lines H1 and HES-2, we found that miRNA-302a, which is highly expressed in undifferentiated cells (Figure 2A, Table S1), can fully repress DsRed even at high copy number (Figure 3B). In contrast, reporters that detect miR-489 and miR-375, miRNAs that are not expressed in undifferentiated hPSC, (Figure 2A, Table S1), only slightly affected DsRed expression (Figure 3B). Our data show that sensors detecting miR-489 and miR-375 are expressed at significantly lower levels than a control construct that does not contain any target sites in DsRed (n.t.) but were significantly higher than constructs that contain mock miRNA target sites FF3 or FF6 (Figure S2E). Thus, we conclude the observed small but significant changes in DsRed expression are due to differences in 3’UTR sequences and miR-489 and miR-375 are, as expected, not substantially active in hPSC. Additionally, we tested the same set of miRNA sensors in hPSC-derived definitive endoderm (DE) and observed significant repression of the miR-375 sensor, but no significant changes in miR-489 and miR-302a sensors (Figure 3B) in agreement with published expression levels (data labelled as Day 2 in Figure 2A, Table S1). Finally, we tested HEK293 cells that show a miRNA expression profile similar to mesenchymal- and neural progenitor cells (MPC and NPC) in our data set (Figure 2A). As expected, the DsRed signal of all three sensors was not substantially repressed in HEK293 (Figures 3B and S2E).

**Figure 3.**
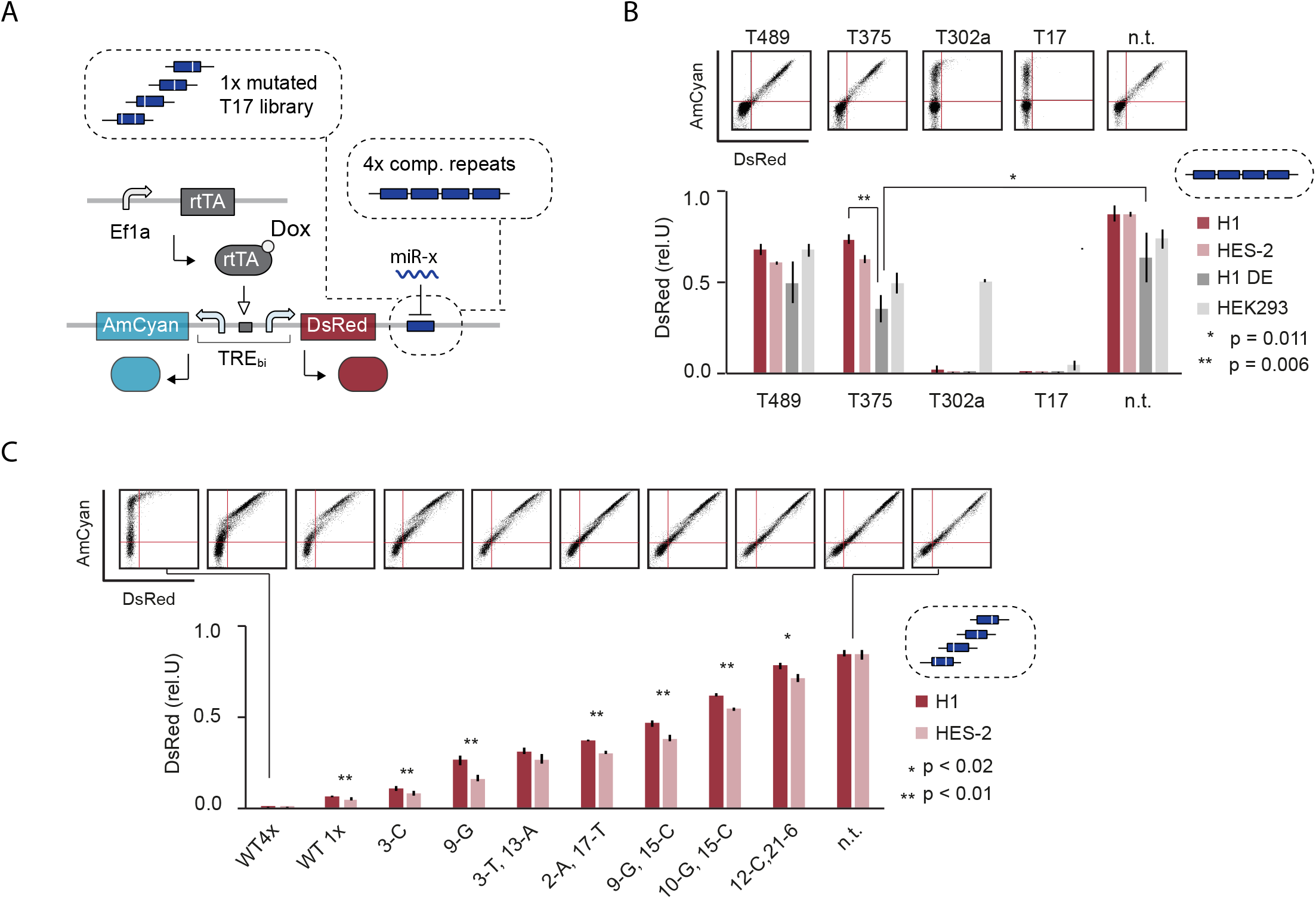
Endogenous miRNAs allow discrete and analog control of synthetic genes in hPSC. **A)** Illustration of bi-directional miRNA sensor system. **B)** Bar chart showing relative DsRed expression of sensors containing the indicated miRNA target sites as four fully complementary repeats (discrete miRNA sensing). The control vector (n.t.) does not contain any target sites. Two-tailed unpaired t-tests were performed to compare the definitive endoderm derived from H1 (H1 DE) to H1 and the control vectors. For comparison to H1, data were normalized with the control vector (nt) before performing the t-test. P values that were equal to or smaller than 0.01 are indicated. Representative scatterplots of HES-2 are shown on top. **C)** Bar chart showing relative DsRed expression of sensors containing different mutated target sequence of miR-17 as listed in Table S4. Additionally, a sensor containing no target (n.t.) and two wildtype (WT) T17 target sites with 1x and 4x repeats were tested. Two-tailed unpaired t-tests were performed to compare H1 and HES-2 for each sensor. P values as indicated. Each bar chart corresponds to mean +/− s.d. from three biological replicates. Experimental description in Table S6. See also Figure S2.

We then used the same bi-directional sensor system to test if synthetic genes can be fine-tuned in hPSC in an analog manner (Figures 3A and 3C). Specifically, we cloned, into the 3’ UTR of DsRed, eight variants selected from previously published miRNA-17-based misFITS library [45], where each variant differs from a perfectly complementary site by one or two nucleotides (Table S4). We also tested 1x and 4x fully complementary miR-17 target sites. Upon transient transfection, we measured DsRed expression relative to untargeted AmCyan in two hPSC lines (HES-2 and H1). We found that each target variant leads to a distinct and defined DsRed expression level spanning the entire dynamic range from fully repressed to fully activated (Figure 3C). Compared to the HES-2 line, the H1 line showed significantly higher DsRed expression for most sensors (Figure 3C).

In summary, our data indicate that miRNA expression data accurately predict relative repression strength of all three computationally identified miRNAs in hPSC, with near discrete sensing behavior. Further we show that endogenous miRNA-17 target libraries allow production of exogenous reporter proteins at pre-specified expression strengths in hPSC thereby enabling analog tuning.

### Model-guided combinatorial screening optimizes circuit performance in hPSC

While miR_low_ sensors typically achieve high dynamic ranges [35, 40, 41], it has been difficult to engineer miR_high_ sensors (here to detect miR-302a) with high dynamic ranges within the full bow-tie configuration [34, 35, 46, 47] (Figure 4A). Reasons include the length of the signaling cascade, the unfavorable dynamics of certain components and substantial variation in expression strength, growth rate and transfection efficiency across different human cell types. Thus, we undertook a model-guided circuit optimization approach with the goal of increasing the On / Off ratio of the miR_high_ sensor within an hPSC embedded bow-tie circuit.

**Figure 4.**
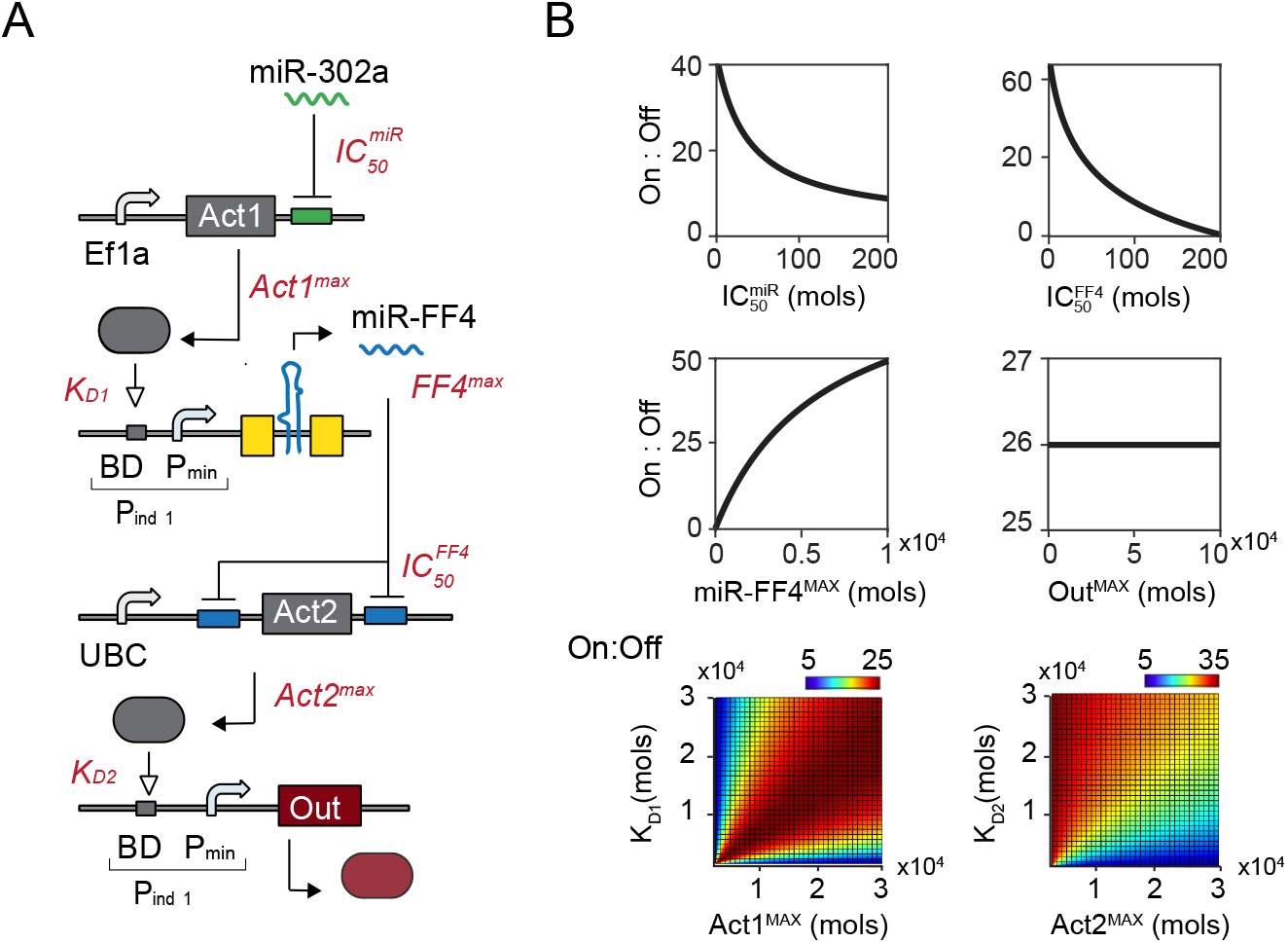
Modeling and parameter screening of miRhigh sensor within whole circuit. **A)** Circuit schematic depicting the miRhigh sensor within the whole bow-tie circuit. The miRhigh sensor recognizes miR-302a in the 3‘ UTR of a constitutively expressed synthetic transactivator (Act1). Act1 induces a synthetic miRNA (miRFF4) that in turn represses a constitutively expressed transactivator Act2. Act2 induces expression of an output protein (here mCherry). In red: key parameters of each reaction. Act1^MAX^ and Act2^MAX^ are the maximal asymptotic expression achieved by the constitutive promoter driving the activator. Out^MAX^ and FF4^MAX^ are the maximal asymptotic expression when the inducible upstream promoter is fully activated. IC50^miR^ and IC50^FF4^ represent miRNA concentrations that lead to 50% knock-down in production of the protein targeted by the given miRNA. Associated equations in Methods. **B)** Dose response curves of individual parameter screens (right, top and middle) and color plots of two-parameter screens (right, bottom) for calculated On / Off ratio. See also Figure S3.

We adapted the mathematical model described in Schreiber et al. [46] to simulate the performance of the miR_high_ sensor within a bow-tie framework (Figure 4A). In line with previous models [39, 46], our model describes miRNA-mediated repression and transcription-factor mediated activation assuming non-cooperative Hill-like relationships between each upstream and downstream component (Figure 4A, equations in Methods). By performing a parameter screen, we found that the majority of parameters, such as IC_50_^miR^, IC_50_^FF4^ and FF4^MAX^ (Figure 4A and Methods), show similar trends as reported in Schreiber et al. who analyzed the miR_high_ sensor behavior on its own [37] (Figure 4B). An interesting new observation arising from our analysis is that K_D_ and Act^MAX^ requirements are different for Act1 and Act2. Act1 performs similarly to the model presented in Schreiber et al, achieving a high On/Off ratio whenever K_D_ or Act1^MAX^ is not particularly low. Act2, however, only provides a high dynamic range for the circuit when Act2^MAX^ is relatively low. This makes intuitive sense, as high Act2^MAX^ levels are more difficult to repress by miR-FF4 (Figures 4B and S3).

Next, we implemented parameters suggested by our model by creating a small library of parts and mammalian Kozak compatible backbones for assembly using the MoClo system [51] (Table S5). Specifically, we chose a strong promoter (Ef1a [52]) driving Act1 to achieve high Act1^MAX^ and FF4^MAX^ (Figures 4 and S3). Second, we flanked Act2 with three fully complementary repeats of the miRFF4 target site (in the 5’UTR and 3’ UTR of Act2) to achieve a low IC_50_ (Figures 4 and S3) [46, 53]. Third, we chose a relatively weak promoter (UBC [52]) to express Act2 to keep Act1 ^MAX^ levels low (Figures 4 and S3).

Because of the distinct but interconnected performance requirements of Act1 and Act2, we first carefully characterized several different Act1 and Act2 constructs (Figure S4), which showed that composing the miR_high_ sensor of an OR gate, detecting miR302a and miR302b, does highly increase the dynamic range of the FF4 expression (Figure S4A). We then set up a combinatorial screen, where we simultaneously changed Act1 and Act2 concentration, which corresponds to parameters Act1^MAX^ and Act2^MAX^ in our model (Figure 5A-D). For this, we first evaluated, in silico, how a combinatorial screen of Act1^MAX^ and Act2^MAX^ affects circuit performance in our model. Specifically, we estimated FF4- and Output production in On- and Off-state and calculated Off / On ratio for FF4 and On / Off ratio for the Output. We found that Act1^MAX^ affects FF4 production independently of Act2^MAX^ with best Off / On performance at low Act1^MAX^ amounts (Figure 5B). The Output production, on the other hand, is affected by both Act1^MAX^ and Act2^MAX^ with best On / Off ratio at intermediate levels of Act1^MAX^ and low levels of Act2^MAX^ (Figure 5B).

**Figure 5.**
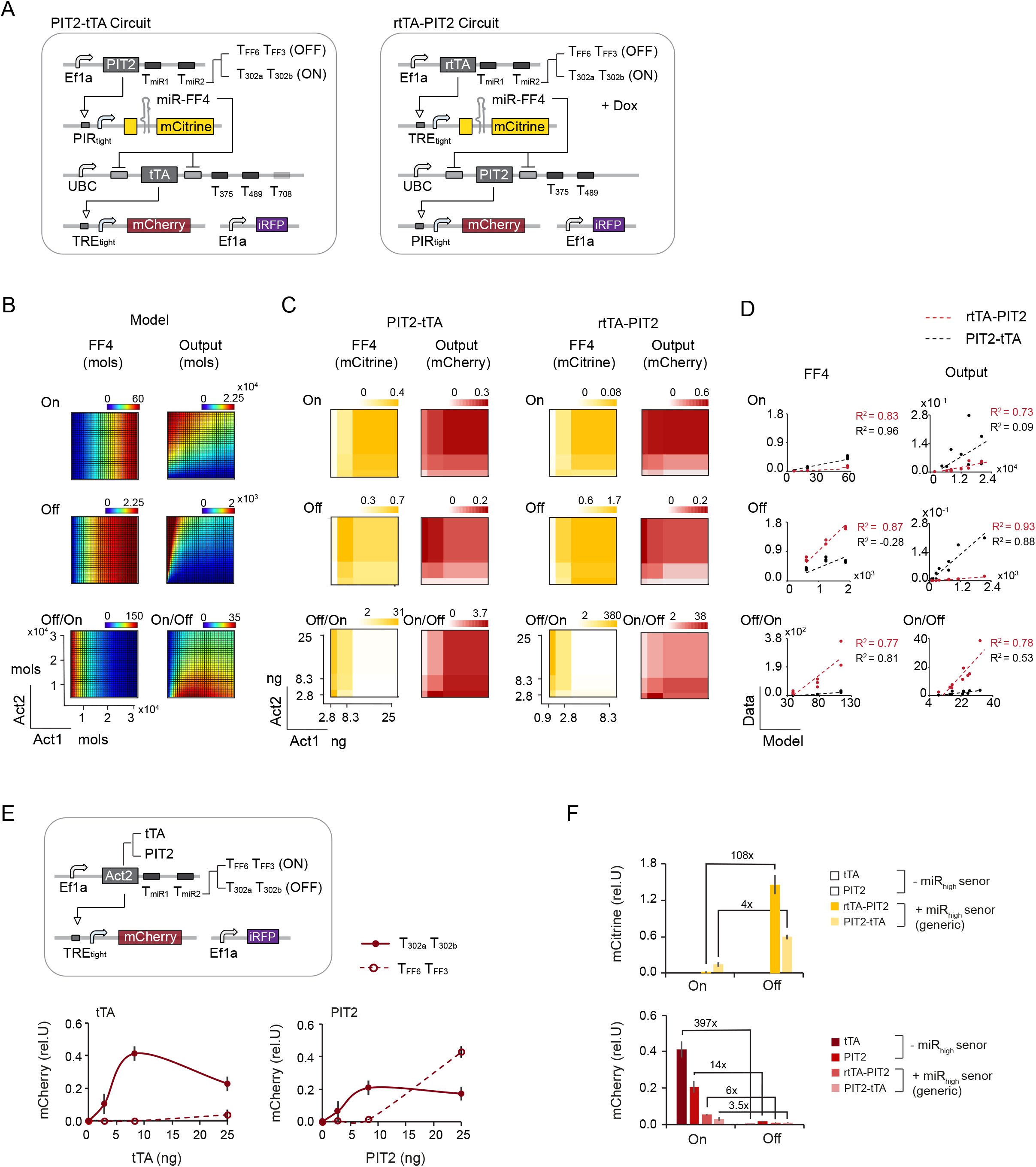
Model-guided combinatorial screening optimizes circuit performance in hPSC. **A)** Circuit schematic. Two different Act1-Act2 circuit configurations are depicted: PIT2-tTA (left) and rtTA-PIT2 (right). **B)** Computational combinatorial screening of Act1^MAX^ and Act2^MAX^ using the model depicted in Figure 4A. Calculated FF4 (left) and Output (right) levels are shown for On, Off and Off/On (FF4) or On/Off (Output). **C)** Experimental combinatorial screening in H1. Amount of Act1 and Act2 expressing plasmids were simultaneously changed at indicated amounts. Color plots show normalized mCitrine (left) and mCherry (right) expression levels for each Act1-Act2 combination. Color bars show lowest and highest expression range on a linear scale with median as the midpoint **D)** Correlation plots with calculated values on X axis (Model) and experimental data on Y axis (Data). A linear regression model (y = a + b*x) was used to compare experimental data with model predictions. Linear fit and coefficient of determination (R^2^) as indicated. PIT2-tTA circuit (black), rtTA-PIT2 circuit (red). **E)** Characterization of circuits operating with miRlow sensors only. Chart shows normalized Output production in response to changing amounts of plasmids encoding Act2, tTA (left) and PIT2 (right). Both transactivators have been tested when targeted by miR302a/b (Off) or by miRFF3/FF6 (ON). **F)** Comparison of the maximal dynamic range across the two generic circuit versions and the two miRlow only circuit versions. For miRlow only circuits and PIT2-tTA circuit, the configuration with highest dynamic range in Output has been selected. For rtTA-PIT2 circuit, because of the low On state in the circuit with highest dynamic range, the circuit configuration with highest Off/On ratio in mCitrine / FF4 and highest On level in mCherry has been selected. Charts and color plots show mean ± s.d. of at least three biological replicates. Experimental description and raw data are provided in Table S6. See also Figure S5.

Experimentally, we tested two circuit configurations with different Act1-Act2 combinations, termed PIT2-tTA- and rtTA-PIT2 circuit (Figure 5A). We combinatorically screened Act1 and Act2, each at three different levels, resulting in nine Act1-Act2 combinations for each circuit. The On state was generated with Act1 containing target sites for miR-302a and miR-302b (T302a T302b) and the Off state with Act1 containing mock target sites (TFF3 TFF6) (Figure 5A). FF4 and Output production were approximated by measuring mCitrine and mCherry expression, respectively (Figure 5C). Because we observed strong inhibition on FF4 / mCitrine expression at high Act1 amounts in the Off state when using the rtTA-PIT2 circuit (Figure S5B), we have used lower amounts of Act1 in the rtTA-PIT2 circuit compared to the PIT2-tTA circuit (Table S6).

To better assess how accurately the data of the two different circuit configurations match our model predictions, we used a linear regression model and calculated the coefficient of determination (R^2^) (Figure 5D). We found that rtTA-PIT2 circuit configuration performance correlates well with model predictions with expected signal transduction properties at all DNA amounts measured (Figures 5D, 5C and S5A). In contrast, PIT2-tTA circuits, some deviated from the model in FF4 / mCitrine expression, caused by inhibition of FF4 / mCitrine expression at high Act1 concentration in the Off state (Figures 5D, S5C), suggesting that lower Act1 levels would have been beneficial for this circuit configuration as well. However, even at lower Act1 levels, Output expression did not perform as expected within the PIT2-rtTA circuit (Figures 5C, 5D and S5A, data labelled with 2.8 and 8.2 ng Act1). Overall, the FF4 / mCitrine levels were markedly lower and mCherry levels markedly higher in the PIT2-tTA compared to the rtTA-PIT2, consistent with observations from individual module testing (Figure S4). We conclude that FF4 production is not high enough in most of the combinations to sufficiently repress Act2 in the PIT2-tTA circuit leading to unexpected circuit behaviour for this configuration (Figures 5C and S5A). Indeed, the best On / Off ratio for the PIT2-tTA circuit was around ∼4-fold, while the best On / Off ratio for the rtTA-PIT2 circuit was around 38-fold. We note, however, that the high On / Off ratio in the rtTA-PIT2 circuit comes at the cost of a very low On state (Figures 5D and S5A). As suggested by the model, the expression levels in the On state can be increased with increasing amounts of Act2 levels with the trade-off to reduce the On / Off ratio (Figures S5A and S6E).

Because a miR_high_ sensor is not always required to discriminate desired cell states, as discussed in Figure 2, we created circuits that operate only with miR_low_ sensors. For this, we tested the Off state with T302a T302b target sites and the On state with TFF4 TFF6 target sites. In this design, we used a Ef1a promoter to drive Act2, because the UBC promoter did not lead to saturation in activation of the downstream mCherry output (Figure S4B). We compared Ef1a-driven tTA with Ef1a-driven PIT2 in H1 and measured the output response to changing activator amounts (Figure 5E). Using this setup, we found that tTA outperforms PIT2, both in terms of dynamic range and the overall repression behaviour in the Off state (Figure 5E). Indeed, the dynamic range of these circuits, with best On / Off ratio at 397-fold and 14-fold for tTA and PIT2, respectively, largely exceeds the performance of the generic circuits that include a miR_high_ sensor (Figure 5F).

To sum up, we used a model-guided combinatorial screen where computationally identified key parameters Act1^MAX^ and Act2^MAX^ were systematically and simultaneously changed within the entire generic circuit. By comparing two different Act1-Act2 circuit configurations, this approach allowed us to quickly identify the circuit and DNA amounts that performed best in hPSCs according to design criteria. Further, we created circuit versions that detect only miR_low_ miRNAs with substantially increased dynamic range compared to generic circuits that operate with a miR_high_ sensor. The here reported designs, plasmid libraries and optimization approach provides a unique foundation for synthetic circuit engineering in hPSC.

### Discrete-to-analog converters allow cell state specific output tuning in hPSC

Having identified a circuit configuration with predictive performance of the miR_high_ sensor, we aimed to assess if our circuit could successfully distinguish between cell states on the basis of endogenous miRNA expression levels. To perform these experiments, we selected the better performing circuit rtTA-PIT2 and tested the circuit in hPSC lines H1 and HES-2 and in HEK293 (Figure 6 A-C). Specifically, we compared the circuit detecting the identified hPSC-specific miRNA input profile (Figure 6A, circuit 1) in comparison to a circuit where T302a / T302b is replaced with TFF3 / TFF6 to measure the maximal FF4 / mCitrine production and Off state (Figure 6A, circuit 2) and a positive control circuit without miR_high_ sensor and mock miR_low_ sensors to measure the maximal Output / mCherry production (Figure 6A, circuit 3). Because HEK293 expresses low levels of miR-302a/b (Figures 6B and S2E), we expected increased FF4 and reduced Output expression in HEK293 compared to hPSC. Indeed, HEK293 expresses FF4 / mCitrine at a very high level, leading to at least 8-fold higher repression of Output / mCherry compared to hPSC lines H1 and HES-2 (Figure 6B, left). Normalizing the data for the observed promoter bias, this results in more than 50-fold output repression in HEK293 compared to hPSC (Figure 6C, right).

**Figure 6.**
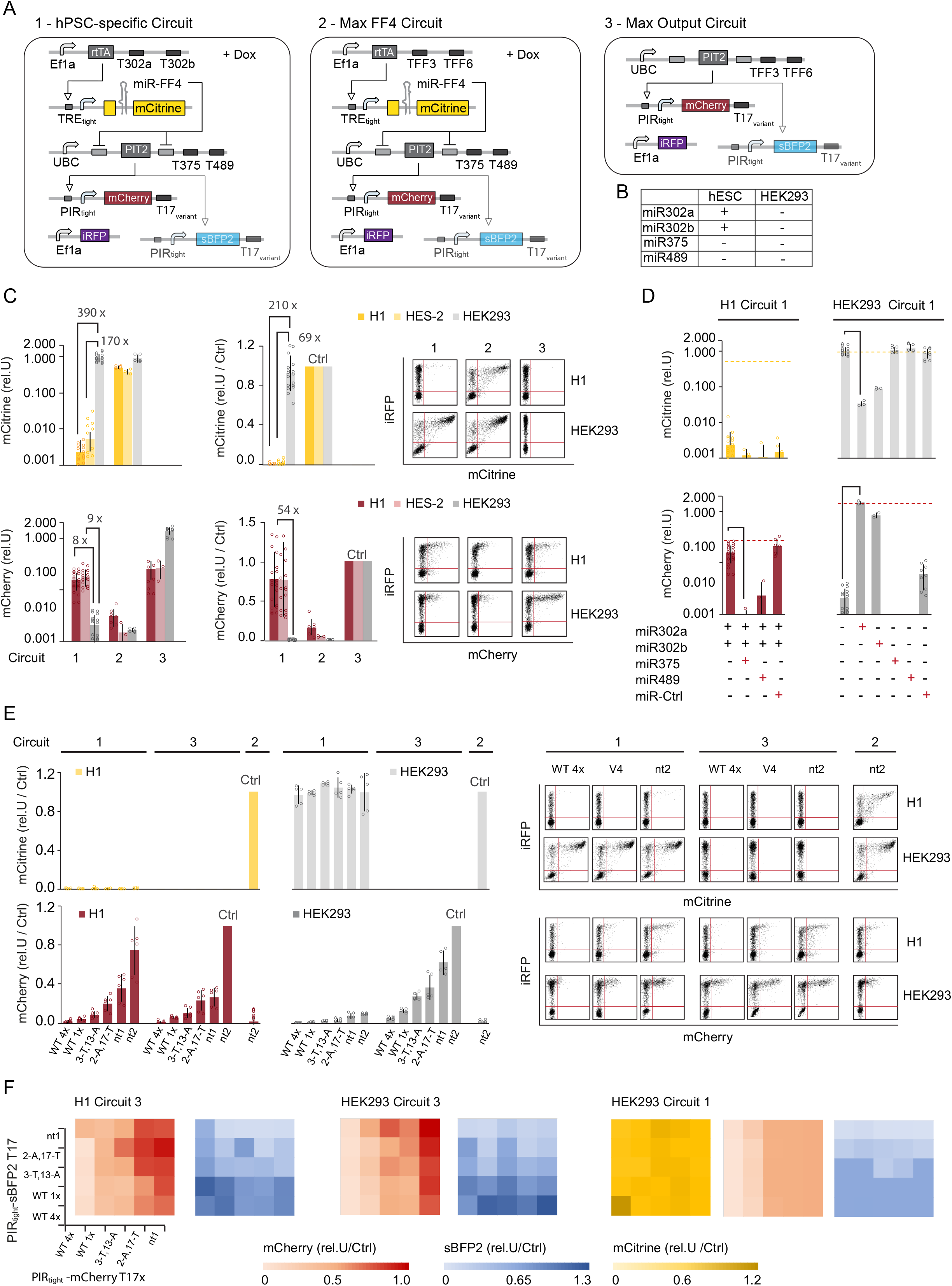
Discrete-to-analog converters allow cell state specific output tuning in hPSC. **A)** Circuit schematics depicting the hPSC specific circuit (circuit 1) recognizing miRNAs miR302a/b, miR375 and miR489 and two control circuits that measure the maximal levels of FF4 / mCitrine (circuit 2) and the maximal output / mCherry level (circuit 3). **B)** Summary of the expression profile for hPSC and HEK293. See also Figure S2E **C)** Bar chart showing the performance of circuit 1 - 3 in H1, HES-2 and HEK293. mCitrine (top) and mCherry (bottom) expression was normalized to internal Ef1a-iRFP control (rel.U.) (left) and further normalized to maximal mCitrine (circuit 2) (top) or maximal mCherry (bottom) (rel.U./Ctrl) (right). **D)** Bar chart showing performance of circuit 1 in response to administration of miRNA mimics in H1 (left) and HEK293 (right). First sample (with profile + + - - - for H1 and - - - - - for HEK292) has no miRNA administered. For the other samples, miRNA dose with highest change in response has been selected from miRNA titration experiments (see Figure S6C), administered miRNA mimic labelled in red. Maximal mCitrine and mCherry levels (as calculated in B) are depicted as yellow and red dashed lines, respectively. Chart is shown in a base 10 logarithmic scale. **E)** Bar chart showing fine tuning of output using miSFITs library. Shown are FF4 / mCitrine (top) and Output / mCherry (bottom) of circuit 1 and 3 for different T17 variants. Data were normalized to control circuits. For C - E, each bar corresponds to mean +/− s.d. from at least three biological replicates. Individual samples are shown as circles. **F)** Color plots showing mean mCherry and mean sBFP2 expression normalized to control circuits. Color plots show mean ± s.d. of at least three biological replicates. Color bars with lowest, median and highest expression values, as indicated. Experimental design and raw data in Table S6. See also Figure S6.

To further characterize circuit performance to changing miRNA amounts and testing of the full logic miRNA input profile, we measured FF4 / mCitrine and Output / mCherry expression in response to externally administered miRNA mimics. Because miRNA and LNA-administration into hPSC has proven difficult using our transfection protocol (Figure S6A-D), we characterized the circuit performance in response to all input miRNAs in HEK293 (Figures 6D and S6C). We found that administration of miR-302a and miR-302b can substantially repress FF4 / mCitrine which results in full release of the Output expression confirming accurate signal processing of the miR_high_ sensor within the bow-tie framework (Figures 6D and S6C). Finally, we found that addition of miR-375 and miR-489 mimics can overcome the remaining leakage in HEK and hPSC (Figures 6D and S6C), suggesting that cell types or cell states that differ in more than one miRNA from the hPSC specific profile will have an even better off target behavior, allowing discrete cell state detection in hPSC.

Confirming the accurate logic input signal processing, we proceeded to evaluate if the output could be fine-tuned within the whole circuit configuration. Because the output level in the On state is relatively low in hPSC, we increased the Act2 amount to 25ng / 96 well leading to a 0.1:1:1:1 molar ratio between the circuit plasmids. This resulted in a 3-fold increase in output level and a 3-fold decrease in On (H1) / Off (HEK293) ratio (Figure S6E). Testing the positive control circuit with a small selection from the miR-17 library, we found that the output could be tuned in hPSC similar as in HEK293 (Figure 6E, circuit 3), with an exception for one variant that behaved different between the two lines (Figure S6F). As expected, we observed reduced output expression in HEK293, but not in H1, when using the PSC specific circuit for all tuners (Figure 6E, circuit 1). Because the highest On state is already weak, by further tuning the levels down, we observed not only a shift in the mean value of the expression intensity but also in the number of cells that scored as mCherry positive (Figure 6E, scatter plots, Table S6).

Finally, we evaluated if we could independently tune two outputs by testing all combinations of miSFITs targeting mCherry and sBFP2 constructs, respectively. When testing circuit 3 in hPSC and HEK293, we found that mCherry and sBFP broadly follow the expected trend from low to high expression for each selected T17 variant (Figure 6F). However, compared to when a single reporter is used, repressive effects are observed when a variant is co-expressed with another highly expressive variant, suggesting competition between PIT2 activators and / or cellular resources [54, 55] (Figure 6F, circuit 3). When testing the hPSC-specific circuit (circuit 1) in HEK293, we found that FF4 / mCitrine is strongly expressed and both outputs are significantly repressed in all output combinations compared to output levels in circuit 3 (Figure 4F, circuit 1), demonstrating that hPSC-specific circuit action can be achieved for all variants in all combinations. Interestingly, we found that the construct without target sites (n.t.1), which is equivalent to the library constructs, showed a significant lower expression strength compared to n.t.2 (Figures 6E and 6F). Constructs n.t.1 and n.t.2 have identical promoter, CDS and polyA sequence but differ in their backbone and UTR sequences (Table S5).

To sum up, our data demonstrate that the hPSC-specific circuits produce substantially higher outputs compared to non-hPSC cells, such as HEK293, and are responsive to all four miRNA inputs. This provides evidence that logic integration of hPSC specific miRNA inputs, approximated with Output = miR-302a OR miR-302b AND NOT miR-375 AND NOT miR-489, is feasible. We also demonstrate that the discrete signaling can easily be converted into analog signaling with the use of miR-17 based miSFITs, allowing the independent fine-tuning of at least two outputs. This first demonstration of a state specific multi-input multi-output system in human cells lays the foundation for numerous applications in regenerative medicine.

### Circuit-mediated BMP4 secretion enables control over cell composition and pattern formation in micropatterned hPSC colonies

Thus far, we have demonstrated the development of a discrete-to-analog converter operating in hPSC. We next set out to deploy these circuits in a proof-of-principle scenario to achieve control over cell fate specification and differentiation outcomes from hPSC. Specifically, we investigated if we could use our circuits to control differentiated (endoderm, ectoderm and mesoderm) cell compositions in responses to endogenously circuit-regulated levels of the developmental morphogen BMP4, which is typically administered exogenously to induce *in vitro* differentiation [56–59].

First, we evaluated our ability to control the secretion levels of BMP4 in the absence of exogenous BMP4 in micropatterned hPSC (RUES2) colonies. For this, we used the circuit version that showed the highest dynamic range in Figure 5, operating without miR_high_ sensor and using tTA as Act2. Act2 controlled a library of miSFITs-targeted BMP4 constructs (Figure 7A). The circuit also controlled iRFP that has not been targeted with the miSFITs library as a way to identify cells that have been transfected. Following our design expectations, we achieved expected graded levels of secreted BMP4 that saturated with the highest expressing tuners (2-A,17-T and 10-G,15-C V4) (Figure 7B, left). iRFP expression, on the other hand, decreased with increasing BMP4 expression, possibly due to cell resource limitations [54, 55] (Figure 7B, right).

**Figure 7.**
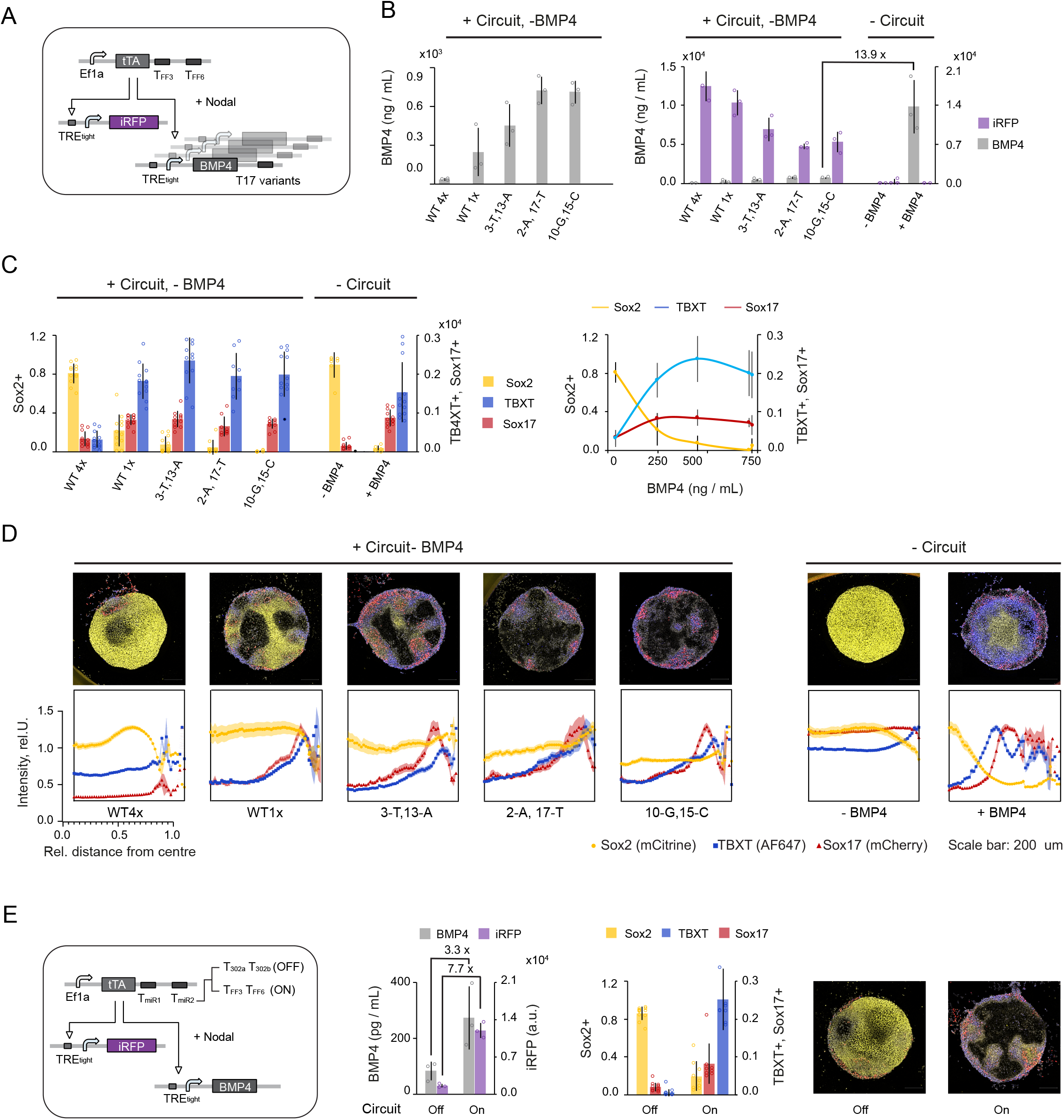
Circuit-mediated BMP4 secretion enables control over cell composition and pattern formation in micropatterned hPSC colonies. **A)** Circuit schematic for fine tuning of BMP4 using miSFITs library. **B)** Circuit-mediated BMP4 secretion using miSFITs library. Micropatterned RUES2 have been transfected with circuit as illustrated in A. Bar chart on the left shows BMP4 concentration for each circuit variant determined from culture media by ELISA. Bar chart on the right shows the same secreted BMP4 concentration along cell internal iRFP expression signal, measured by Flow Cytometry in comparison to untransfected controls in the absence (-BMP4) and presence (+ BMP4) of BMP4 (right). **C)** Bar chart showing the relative fraction of cells expressing a given germ-layer marker as analyzed from confocal microscopy images (right). Dose response chart of the marker fraction in response to measured BMP4 concentration (left). See also representative Flow Cytometry scatter plots in Figure S7B. **D)** Radial marker profiles of each germ-layer marker across all BMP4 conditions analyzed from confocal microscopy images. Charts on the left show normalized intensity of marker expression at a given colony position, where x=0 is the center of the colony and x=1 is the edge of the colony. Representative microscopy image of a single colony (top). **E)** Control of BMP4 in response to the endogenous miRNAs. Micropatterned RUES2 have been transfected with two circuits where tTA contains miR-302a/b target sites (Off) or mock target sites TFF3/TFF6 (On). Each target site is composed of four fully complementary repeats. tTA inducible iRFP and BMP4 have been used without miSFITs targets. Bar chart showing BMP4 concentration as determined by ELISA and average iRFP expression as measured by Flow Cytometry (left). Corresponding bar chart showing the average fraction of cells expressing a given germ-layer marker. Each bar chart show mean ± s.d. from at least three biological replicates for BMP4 and iRFP data and from at least 20 colonies for germ-layer data. Microscopy images in D and E show composite of Sox2/mCitrine (yellow), Sox17/mCherry (red) and TBXT/AF647 (blue). Scale bar represents 200 μm. Raw data and experimental setup in Table S6. See also Figure S7.

Next, we compared how circuit-controlled endogenous BMP4 versus exogenously-supplemented BMP4 regulated differentiation of hPSC colonies. We also tested whether different miSFITs variants, which change BMP4 secretion levels, could be used to control the relative composition of differentiated germ-layer cells. We found that the cells adopted specific fates in a BMP4 dose-dependent manner, where proportion of ectoderm-like Sox2+ cells continually and rapidly decreased and endoderm-like Sox17+ and mesoderm-like TBXT+ cells continually increased until all marker levels were saturated as BMP4 levels saturated (Figure 7C). In terms of cell composition of germ layers, the variant with the lowest BMP4 level (WT4x) resembled untransfected controls in the absence of exogenous BMP4 (-BMP4) while variants with medium to high BMP4 levels resembled colonies with externally-administered BMP4 (+BMP4) (Figure 7C left). Interestingly, the apparent concentration of BMP4 required to achieve the same cell composition was at last 10 times lower when produced endogenously from circuit constructs, compared to exogenous administration (Figure 7B,C).

To evaluate if circuit-controlled BMP4 could induce gastrulation-like pattern formation, we characterized the spatial marker expression from the colonies, and compared the results to externally administered BMP4. For the latter, as previously reported, we observed highly reproducible gastrulation-like patterns [56–59] which are concentric rings of germ layer markers with extra-embryonic-like tissue (CDX2+) at the colony edge (not measured here) followed by Sox17+, TBXT+ and Sox2+ cells radially distributed inwards (Figure 7D, right). In contrast, colonies that expressed BMP4 endogenously from our circuit constructs appeared more heterogenous and showed several sparsely populated regions that did not express any of the three germ-layer markers (Figures 7D, left). Interestingly, BMP4 producing cells, as measured with iRFP, did not retain Sox2 expression when BMP4 was produced at medium to high levels and did not adopt an endoderm (SOX17+) or mesoderm (TBXT+) fate independent of the BMP4 concentration (Figure S7B). Notably, despite the sparsely populated regions, colony-wide patterns formed, with Sox17 and TBXT showing higher expression at the edge of the colony compared to the center (Figure 7D, bottom). The relative radial position of Sox17 and TBXT switched compared to the exogenous BMP4 addition, with Sox17 closer to the center than TBXT, and had become more separated with increasing BMP4 production (Figures 7D and S7C). The qualitatively and quantitatively different pattern formations observed were reinforced by the significantly different AUC measurements for the circuits that expressed BMP4 at low levels versus high levels for the TBXT and SOX17 radial expression profiles (Figures S7D).

Finally, we aimed to show that BMP4 production could be restricted to desired cell states in a discrete manner in response to endogenous miRNA inputs. To demonstrate this, we simulated the Off state with a circuit targeted by miR302a / b and the On state with the mock miRFF3/FF6 target sites. We found that secretion of BMP4 was repressed 3.3-fold when using miR302a//b targets (Off) compared to miRFF3/FF6 targets (On), whereas intracellular iRFP expression showed a 7.7-fold change in On / Off ratio (Figure 7B, left). The dynamic range could be further increased when less tTA construct was used with the trade-off of achieving a lower On state (Figure S7E). Analyzing the number of cells expressing a given germ-layer marker, we found that the Off state, by repressing BMP4, locks the cells into a primarily Sox2 positive state (Figure 7E, right). In the On, around 10% of cells express Sox17 and 10% TBXT, while the fraction of cells expressing Sox2 is less than 30%, indicating the differentiation into the three germ-layers (Figure 7E, right).

In summary, our data indicate that the minimal version of a discrete-to-analog circuit can create different germ-layer compositions from hPSC in a fine-tuned manner with consistent BMP4 dose-dependent cell fate changes. Additionally, BMP4 production can be effectively repressed in response to cell state specific miRNAs, providing evidence that cell state specific analog control of secreting factors can be applied for cell composition control in hPSC. Further, we showed that recombinantly released BMP4 can trigger pattern formation in micropatterned hPSC colonies that are distinct from patterns previously reported using exogenous BMP4.

## Discussion

Cells constantly process signaling inputs in a cell state and context specific manner. Despite impressive achievements in the field of synthetic biology, rational engineering of artificial circuits that better represent the capabilities and complexity of natural biological computing systems remains challenging. Here we demonstrate that discrete-to-analog signal conversion, a form of digital-to-analog conversion [60, 61], can be exploited to exert precise cell state specific control of desired proteins in hPSC. Specifically, we have implemented gene circuits that detect a hPSC-specific miRNA profile and can produce up to two protein outputs that can be individually tuned to desired levels. Our circuits merge two critical functionalities: first, they allow restriction of circuit action to desired cell states, which is crucial for applications where conditional control of desired biomolecules is required; and second, they can produce the desired biomolecule at defined doses, which is important in many systems where the test biomolecule exerts its function at precise physiological ranges. These functionalities can be useful for various applications but fit particularly well with the challenges of controlling developmental processes. During development, different intermediate cell states or lineages differentially control a small set of effectors or regulatory factors that operate at defined amounts. This happens through very complex, often poorly understood, interactions between gene regulatory networks (GRN) and environmental signals [3]. Here we show that the same function can be achieved with a rationally engineered, synthetic and programmable system that operates orthogonally in hPSC and during early differentiation, opening the door to control the production and dose of desired natural or synthetic factors from desired cell states in emerging hPSC-derived 2D and 3D tissues.

Classification of cell states and circuit design are non-trivial tasks. Here we have employed a previously-established computational algorithm that automates the classification and circuit design procedure [39]. While the previous study examined the theoretical performance of this toolbox in the context of cancer classification [39], we have demonstrated and expanded its usefulness in the context of stem cells and changing states during differentiation. Interestingly it appears that undifferentiated hPSC can be discriminated from different cell states with only a few miRNAs. We have further demonstrated that a deterministic biochemical model can give useful and unexpected insights into the performance of a circuit of this complexity, and can be helpful for reducing the number of genetic parts that need to be tested, a significant advantage for applications in hPSCs. Interestingly, the better performing circuit (rtTA-PIT2 circuit) used the same transactivator combination as used in our previous bow-tie implementation created in Hela cells [34]. Because many differences exist between the pervious HeLa and this hPSC circuit implementation, such as a different version of miR_high_ sensor (here adopted from Schreiber et al [46]), different miRNAs inputs and different vector backbones, it suggests that the correct combination and levels of synthetic transactivators (Act1 and Act2) seem to be a main driver of optimal circuit performance and cannot easily be exchanged within the circuit architecture. It further suggests that the other parameters within the circuit have been sufficiently optimized within previous implementations [34, 35, 46]. We also showed that creating a circuit version without miR_high_ sensors is dramatically less time consuming and leads to a higher dynamic range.

Further, our work demonstrated that the use of miSFITs [45] is a straightforward approach to achieve defined analog responses in hPSC. In fact, we observed high dynamic ranges using a bi-directional reporter system in multiple hPSC lines. We further documented that a few of the mutated T17 target sites behave differently in the context of a different 3’UTR and between different cell lines (hPSC vs HEK293), consistent with previous observations [45]. This suggests that either prior selection through screening or a comprehensive optimization of 3’UTR and backbones to enable consistent behaviour across different cell types might be beneficial for certain applications. Importantly, we also demonstrate that miSFITs are an excellent option to tune multiple proteins independently in the same cell or sample. In fact, within our architecture this would not have been possible with inducible systems that tune outputs using externally administered chemical inducers (such as pristinamycin for PIT2 or doxycycline for rtTA) [62, 63]. Applying miSFITs for fine tuning also renders the system suitable for *in vivo* applications where chemical inducers are difficult to administer at defined levels, and actuation is temporarily limited. Other means to change the dose, such as changing the transcriptional activity by modifying the Act2-inducible promoter is possible but inherently more difficult because it will generate a different dose response function for output production, affecting the overall logic sensing performance and with this the entire system. Our data also revealed that precise tuning of two or more output genes using plasmid transfection is challenging because of cellular and circuit-based resource competition. Recent studies characterized how synthetic genes compete with each other for cellular resources in human cells, with computational models that capture and practical solutions to dampen this competition effect [54, 55, 64]. Future efforts to mitigate, or accurately predict cellular resource competition will help improve overall circuit performance and the utility of miSFITs for tuning multiple genes independently.

Finally, we demonstrated that physiological factors such as BMP4 can be secreted into the cultivation media and fine-tuned with miSFITs technology, allowing us to control differentiated cell composition. Although transient transfection and the relatively low number of output-producing cells may be a disadvantage for certain applications, we show that our setup is suitable for applications that aim to control secreted physiological factors, where the effect can be propagated across the population level. Further, we took advantage of context explorer [65] to rapidly quantify spatial marker expression in individual colonies and found that colony-wide pattern formation could be achieved with circuit induced BMP4, despite short range diffusible effects and unequal cell density distribution, highlighting the strong self-organization potential of hPSC. Interestingly, circuit-released BMP4 achieved differentiation and pattern formation at substantially lower concentrations compared to exogenously applied BMP4, creating opportunities to generate desired cell types in a manner that may more closely resemble natural developmental processes [66].

Moving forward, the computational and genetic tools developed here can be applied to generate new customized circuits that control, not only desired secreted factors, but also transcription factors or small regulatory RNAs, from desired cell states or developmental stages. Our circuits in combination with micropatterning, gastruloid or organoid technologies open the door to gain new insights into molecular mechanisms that guide pattern formation and other fundamental processes during early development or organogenesis. Additionally, our circuits can be applied for targeted killing of undesired cell types during differentiation, cell state tracking, re-enforcement (or lock-in) of desired cell states, direct conditional cell reprogramming *in vitro* and *in vivo* and for conditional production of therapeutic agents from hPSC-derived cell products or *in vivo*.

## Limitations

There are several key limitations that need to be addressed before discrete-to-analog converters can become widely applicable to program hPSC, hPSC-derived products and other human cells. First, the current system works robustly with a high dynamic range only when miR_low_ sensors are used, but produces very low yield when the input module comprises a miR_high_ sensor. Thus, the On state and dynamic range of the circuit output needs to be further increased when miR_high_ sensors are required for cell state identification. This could be achieved with extended combinatorial screening using an extended library of genetic parts [67]. Further improvements can be achieved by optimizing the dynamics of the parts, such as delaying the production of Act2 with recombinase-based systems, which can remove leakage in transient systems [9, 47]. Second, due to the long cascade of transcriptional components, the circuit is relatively slow in responding to changes. Thus, implementations of a circuit that aims to discriminate very close states, such as hPSC and DE in our case, may require a faster turnover rate of the circuit proteins and / or stable integration of the circuits. Alternatively, faster acting RNA-only [68] or protein-only [25] circuits could be merged with miSFITs technology. Third, the large circuit size limits their utility for viral-based delivery strategies and with this their *in vivo* applications. Generally, stable integration of circuits of this complexity have proven challenging due to the low integration efficiency in hPSC and because, once integrated, circuit silencing and unexpected cross-activation from genome to the circuit or within circuit components can occur [14, 69]. With current fast development of genome editing tools, automated procedures and new gene regulatory tools for mammalian cells, we believe these challenges can be overcome in the near future to enable straightforward use of digital-to-analog converters for advanced differentiation control of hPSC, direct reprogramming, basic research and beyond.

## Materials and Methods

### Automated miRNA identification

miRNA expression data were obtained from published sources here referred as Bar et al [50], Lipchina et al [49] and Fogel et al [48]. Cell source and miRNA expression data are summarized in Table S1. miRNA Sequencing data from Fogel et al, were renamed according to miRBase nomenclature and mean values of replicates were calculated (Tables S1). In order to access miRNA expression data from different published sources that use different miRNA naming conventions, we modified the pre-processing pipeline of the code developed by Mohammadi et al. [39]. Instead of utilizing the miRNA names as provided by the sources, the provided miRNA sequences were determined by referring to the miRBase versions specified by the source papers. Furthermore, hairpin miRNA sequences were also obtained from the respective miRBase versions, and they were matched and eliminated from the dataset. Then, miRNA names, sequences, and sequencing data from the three sources were compared and combined to generate a merged dataset. Lastly, similar miRNAs within this merged dataset were combined just as Mohammadi et al. did for a single dataset. The optimal input miRNA set was searched based on this finalized merged dataset.

The algorithm with the modified pre-processing pipeline can be accessed on GitHub: https://github.com/jwon0408/SynNetModified.

To run the algorithm, total maximal miRNA inputs and the maximal miRNA input number of each gate have been modified as shown in Tables S2 and S3 (constraints tab). Raw output data files for Figure 2 and Figure S2 can be found in Table S2 and S3.

### Modeling and Simulation

Circuit modeling and simulations were performed in MATLAB. The model describes a miR_high_ sensor within a bow-tie architecture detecting one miRNA-input (miR-302a) and controlling one protein output (as depicted in Figure 4A). The model was expanded from Schreiber et al [46] and assumes non-cooperative Hill-like relationships for miRNA-mediated repression and transcription-factor mediated activation.

The following four steady state equations were used to describe the circuit reactions: miRNA-mediated Act1 repression (1), Act1-mediated FF4 production (2), miR-FF4-mediated Act2 repression (3) and Act2-mediated output production (4):

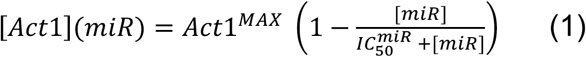

Where [Act1] is the calculated steady-state concentration of the transactivator; Act1^MAX^ is the maximal steady-state activator concentration, without any miRNA mediated repression. [miR] stands for miRNA input concentration and for input miRNA concentration that elicits half the knock down.

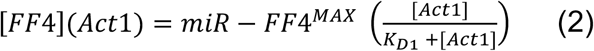

Where [miR-FF4] represents steady-state concentration of miR-FF4, [Act1] is the activator level computed with equation (1), *K*_D1_ is the dissociation constant of Act1 from its promoter. miR-FF4^MAX^ is the maximal expression level of miR-FF4 from an inducible promoter under transactivator saturation.

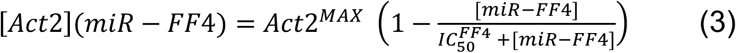

Where [Act2] is the steady state concentration of the central knot activator; Act2^MAX^ is the maximal steady-state activator concentration without any miRNA mediated repression. [miR-FF4] is the miR-FF4 concentration calculated in equation (2) and is the miR-FF4 concentration that elicits half the repression.

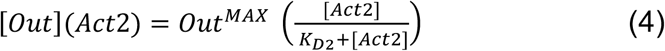

[Out] is the steady state concentration of the circuit output and Out^MAX^ is the maximal output concentration from an inducible promoter under Act2 saturation. [Act2] is the activator concentration calculated in (3). *K*_D2_ is the dissociation constant of Act1 from its promoter.

The parameters for numerical simulations (Figure 4A and S3) were derived from Mohammadi et al [39]. Specifically, we used their optimized parameter set: 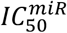: 20 molecules/cell, 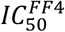: 20 molecules/cell, *K*_D1 =_ *K*_D2_ : 10,251 molecules/cell, Act1^MAX^ = Act2^MAX^: 9,755 molecules/cell, miR-FF4^MAX^: 3,000 molecules/cell, and Out^MAX^ 30,000 molecules/cell. The ON state was simulated with [miR] = 3000 molecules/cell and the Off state with [miR] = 0 molecules/cell. For each parameter scan the varied parameters were tested in the range indicated in the plot and the fixed parameters were set at the numbers indicated above. For the correlation plot in Figure 5D we used 30’000, 10’000 and 3333 molecules per cell for both Act1^MAX^ and Act2^MAX^ to calculate FF4 and output production.

### Cloning

Bi-directional miRNA sensor plasmids were cloned using standard cloning techniques. Briefly, different miRNA target sites (Table S5) were designed with SalI and NotI overhangs, ordered as ssDNA from Sigma Aldrich, annealed by temperature decrease from 95 °C to 4 °C and phosphorylated using T4 Polynucleotide Kinase (NEB, M0201) according to manufactures’ protocol. Annealed fragments were purified using the GenEluteTM PCR Clean-Up Kit (Sigma Aldrich, NA1020). Bi-directional precursor construct (pZ073) [35] obtained from Benenson lab at ETH Zurich was digested with 30 U of SalI-HF (New England Biolabs (NEB), R3138S) and 30 U of NotI-HF (NEB, R3189) using CutSmart Buffer in a total reaction volume of 50 µL for fours at 37 °C. Digested backbone was purified from 0.7 % Agarose Gel using the GenEluteTM Gel extraction Kit (Sigma Aldrich, NA1111). Vector and miRNA target inserts were ligated at 1:10 molar ratio using 400 U of T4 DNA Ligase (NEB, M0202) using provided Ligase buffer in a reaction volume of 20 µL over night at 16 °C. Then 4 µL of ligation mix was transformed into 50 µL of home-made chemically competent *E.coli* DH5alpha obtained from New England Biolabs (C2987I) and selected on LB Agar plates containing 100 µg/mL Ampicillin (Sigma-Aldrich, A0166-25G)

All remaining plasmids were generated using Goldengate cloning with the MoClo toolkit (Addgene Kit#1000000044) as described in Weber et al. [51]. Briefly, a level 0 constructs were generated by cloning a given part (promoter, miRNA target sites, spacer sequences, protein coding sequences or polyA) into level 0 destination vectors from the MoClo toolkit or into home-made level 0 vectors that can restore the Kozak sequence (Table S5). The DNA parts were either I) ordered as dsDNA from Twist Bioscience, II) as ssDNA oligos from IDT Technologies or Sigma Aldrich, annealed and phosphorylated as described above or III) PCR amplified from other vectors using Phusion® DNA polymerase (NEB, M0530L) according to manufactures’ protocol. PCR fragments were analyzed using Agarose Gel Electrophoresis and purified from Gel using the GenElute Gel Extraction Kit (Sigma, NA1111) or MinElute Gel Extraction Kit (Qiagen, 28604) according to manufactures’ protocol. Cloning of level 0 vectors was performed with 40 fmol of level 0 backbone and 40 fmol of DNA insert using 10 U *BpiI* (Thermo Scientific, ER1011), 400U T4 Ligase (NEB, M0202S), 0.15 µL BSA (NEB, B9000S) and 1.5 µL T4 ligation buffer (NEB, B0202S) at a final volume of 15µL. Assembly was performed using the following cycle condition: 37 °C 15 min (1 cycle), 37 °C 2 min followed by 16 °C 5 min (50 cycles), 37 °C 15min, 50 °C 5 min, 80 °C 5 min, 4 °C until use. Then 4 µL of the MoClo mix was transformed into 50 µL NEB 5-alpha competent *E. Coli* (NEB, C2987I) or into 50 – 100 µL home-made chemically competent E.coli 5-alpha. Clones were selected with *lacZ* based blue-white screening method on LB agar plates containing 100 µg/mL Ampicillin (Sigma-Aldrich, A0166-25G), 100 µM IPTG (Sigma-Aldrich, I6758-5G) and 40 µg/mL X-gal (BioShop Canada, XGA001.1).

Expression units were assembled from cloned level 0 parts into level 1 destination vectors provided by the MoClo kit using 40fmol of each vector and 20 U of *BsaI-HFv2* (NEB, R3733L) with remaining conditions as described above. Clones were selected based on the same *lacZ* selection strategy as described above but with 50 µg/mL Spectinomycin (BioShop Canada, SPE201.5) instead of Ampicillin for selection.

All vectors used for assembly and transfections were purified with GeneJET plasmid miniprep kit (Thermo Fisher Scientific, K0503) or GenElute^TM^ Plasmid MiniPrep Kit (Sigma, PLN70). Sequence of every cloned plasmid was verified using Sanger Sequencing performed at the Centre for Applied Genomics at the Hospital for Sick Children, University of Toronto or at Genewiz Inc. Prior to transfections, plasmid quantity was determined using Nanodrop and quality was verified on 0.6-1% Agarose Gels. A complete list of plasmids cloned in this study can be found in Table S5. Constructs that are not restricted for sharing under MTA are provided on Addgene as indicated in Table S5.

### siRNA, miRNA mimics and miRNA inhibitors

MiRNA mimics were obtained from Dharmacon as Human miRIDIAN microRNA Human mimics. hsa-miR-302a-3p, (C-300653-05-0002), hsa-miRNA-302b-3p (C-300669-05-0002), has-miR-489-3p (C-300749-07-0002), hsa-miR-375 (C-300682-05-0002) and miR-Ctrl Negative Ctrl #1, (CN-001000-01-05). siRNA FF4 was custom designed as Silencer^TM^ Select siRNA from Thermo Fisher Scientific (see [27] for sequence) and compared to siRNA Ctrl: Silencer Select Negative Control No. 1 siRNA (Thermo Fisher Scientific, 4390843). miRNA inhibitors were purchased from Qiagen as miRCURY LNA miRNA inhibitors: LNA-miR-302a-3p (YI04100713-ACA), LNA-miR302b-3p (YI04101540-ACA) and used in comparison to the Negative Control B LNA (YI00199007-ADA).

### Cell culture

hPSC lines H1 (WAe001-A) [70] and HES-2 (ESIBIe002-A) [71] were obtained from Christina Nostro and Gordon Keller lab at the University of Toronto, respectively. H1 and HES-2 were thawed and maintained on irradiated mouse embryonic fibroblasts (MEF) and cultured for 6 days in maintenance medium comprising Dulbecco’s minimum essential media DMEM/F12 (77.5 % v/v, Gibco), Knockout Serum Replacement (20 % v/v, KOSR, Gibco), GlutaMax^TM^ (2 mM, Invitrogen; 35050061), penicillin/streptomycin (0.5 % v/v, Gibco; 15140122), non-essential amino acids (1% v/v, Gibco), β-mercaptoethanol (0.1 mM, Sigma) and 20 ng/mL basic fibroblast growth factor (bFGF; PeproTech). Cells were maintained at 37 °C humidified air with 5 % CO_2_ with daily medium exchange. Before transfection, cells were transferred and grown in Essential 8^TM^ Medium (Gibco, A1517001) supplemented with 0.5 % Penicillin/Streptomycin Solution (Gibco; 15140122) on 2 % Geltrex^TM^ (Gibco, A1413302) coated plates at a split ratio of 1:6 for at least one passage. Media was exchanged daily and splitting was performed every 4-5 days when cells reached 75-85% confluency using TrypLE^TM^ (Gibco, 12605028). Cultures were propagated for at most 4 passages before being replaced by fresh cell stock.

The hPSC RUES2 line [72] was obtained from Prof Ali Brivanlou’s lab at the Rockefeller University. The cell line was cultured using Geltrex (Life Technologies, Catalog # A1413301, 1:50 dilution) and mTeSR medium (StemCell Technologies, Catalog # 85850) without penicillin/streptomycin. Cells were clump passaged using ReLeSR (Stem Cell Technologies, Catalog # 05872) and maintained at 37 °C with 5 % CO_2_. Media was changed daily, and cells were passaged once they reached 75-80% confluence.

HEK293 cells were obtained from Princess Margaret University Health Network (Pan’s lab) and cultured at 37 °C, 5 % CO_2_ in DMEM (Gibco; 11965092), supplemented with 10 % fetal bovine serum (FBS; Gibco; 12483020), 1 % GlutaMAX^TM^ (Gibco; 35050061) and 1% Penicillin/Streptomycin Solution (Gibco; 15140122). Splitting was performed using 0.25% Trypsin-EDTA (Gibco; 25200072) every 2–3 days until cells reached 90-100 % confluency.

All cell lines have been tested for Mycoplasma at the Hospital for Sick Children. hESC H1 and HES-2 have been karyotyped at Thermo Fisher Scientific and WiCell.

### Transfections

All transfections of hESC H1 and HES-2 were performed using Lipofectamine Stem Transfection Reagent (Thermo Fisher Scientific, STEM00015). For transfections in pluripotency state, cells were dissociated into single cells with TrypLE^TM^ for 4-5 minutes at 37 C and seeded into 2 % Geltrex coated 96- or 12-well plates (Fisher Scientific) in Essential 8^TM^ Medium (Gibco, A1517001) supplemented with 0.5 % Penicillin/Streptomycin Solution (Pen/Strep, Invitrogen; 15140122) and x 10 µM Y27632 Rho-rock inhibitor (Reagents Direct, 53-B85). 1-1.2 x 10^4^ cells were seeded into 100 µL culture media per 96-well for transfections without antibody staining and 1 x 10^5^ cells were seeded into 500 µL culture media per 12-well for transfections with subsequent antibody staining. Cells were incubated at 37 °C, 5% CO_2_ for around 24h until transfection was performed at around 55-65% confluency. For transfections with subsequent differentiation into endoderm lineage, cells were seeded as patches into 12-well plates on 2 % Geltrex using NutriStem® hESC XF (Biological Industries, 05-100-1A) supplemented with 0.5 % Penicillin/Streptomycin Solution. Cells were grown for two to three days before transfection was performed at around 90% confluency. At the day of transfection, medium was replaced with fresh growth medium (Essential 8 or Nutristem respectively) supplemented with Doxycycline hyclate (Dox, Sigma, D9891) at a final concentration of 0.8 μg/mL if required. Plasmid DNA and miRNA mimics, if needed, were mixed according to Table S5. Lipofectamine Stem Reagent was diluted 25-fold with Opti-MEM I Reduced Serum (Life Technologies 31985-962). For each 96 well and 12 well sample, 10 µL and 100 µL respectively of diluted reagent was mixed with the DNA mixture, incubated for 10-12 minutes at room temperature and added to the cells. For samples in pluripotency state, medium was replaced after 24h with fresh Essential 8^TM^ including Pen/Strep and Dox if needed but no Rho-Rock inhibitor using a volume of 150 µL and 1.5 mL per 96-well and 12-well, respectively. Cells were harvested for flow cytometry and antibody staining 48 h after transfection. For differentiations, medium was replaced 24 h after transfection with endoderm inducing media (see below).

### Endoderm Differentiations

hESC H1 cells were seeded from MEF condition into 12-well plates and grown in Nutristem hESC XF (Biological Industries, 05-100-1A) on 2 % Geltrex for 1-2 days until they reached 90 % confluency. Some samples were transfected using Lipofectamine Stem Reagent as described above, untransfected samples were kept in fresh Nutristem. 24h after transfection (=Day0), endoderm differentiation was initiated. At day 0, pluripotency medium was replaced with endoderm base medium (RPMI1640 (Gibco, 11875093), 1 % GlutaMAX^TM^, 1 % penicillin-streptomycin solution) supplemented with 2 µM CHIR 99021 (Reagent Direct, 27-H76) and 100 ng/mL Activin A (home-made). After 24 h, medium was replaced with endoderm base medium supplemented with 100 ng/mL Activin A, 50 µg/mL Ascorbic Acid (Sigma-Aldrich; A8960) and 5 ng/mL human FGFb (Peprotech; 100-18B). Medium was replaced daily for 3 days using 1.5 mL per 12 well. On day 2 after endoderm induction, medium was supplemented with Doxycycline at a final concentration of 0.8 μg/mL for samples transfected with bidirectional reporter systems and some of the control samples.

### Micropatterned plate preparation

The protocol employed has been reported elsewhere [73, 74]. Briefly, custom Nexterion-D Borosilicate (Schott, D263) thin glass coverslips (dimensions 110×74 mm) were spin-coated with Lipidure-CM5206^TM^ (Lipidure) (NOF America Corporation). Lipidure was reconstituted in 100% ethanol at 2.5 mg/mL. The coverslips were sterilized by covering their side with isopropanol alcohol and spinning them at 2500 rpm for 30 s on the Laurell Spin Processor (Laurell Technologies Corporation, Model # WS-650Mz-23NPPB). This was followed by addition of 1.5 mL Lipidure solution on the sterilized side of the coverslip and spinning it at 2500 rpm for another 30 s. To create micropatterns, the Lipidure coated side of the coverslip was exposed to deep UV for 20 min through a quartz photomask. The diameter of each micropattern was 1 mm. The coverslips were subsequently attached to 96-well bottomless microtiter plates (VWR, Catalog # 82050-714) using epoxy (Loctite, M-31CL). Carboxyl groups on the photo-activated regions were activated with 50 µL/well of 0.05 g/mL N-(3-Dimethylaminopropyl)-N’-ethylcarbodiimide hydrochloride (Sigma-Aldrich, 03450) and N-Hydrosuccinimide (Sigma-Aldrich, 130672) solution for 20 minutes. The wells were then rinsed followed by the addition of 2 % Geltrex. The plate was subsequently left on an orbital shaker overnight at 4 °C.

### BMP4-dependent differentiation of micropatterned RUES2 colonies

Micropatterned plates were thoroughly rinsed before the addition of cells. RUES2 cell suspension was created using TrypLE when the culture had reached 75-80 % confluence. Cells were seeded at a density of 25,000 cells/well with the seeding medium containing the ROCK-inhibitor (ROCK-I) Y-27632 (Reagents Direct, 53-B85-100). The seeding media consisted of DMEM, Knockout Serum Replacement (ThermoFisher, 10828028), Penicillin/Streptomycin (ThermoFisher, 15140122), 2-Mercaptoethanol (ThermoFisher, 21985023), Glutamax (ThermoFisher, 35050061), Non-Essential Amino Acids (ThermoFisher, 11140050), B27 without Retinoic Acid (ThermoFisher, 12587010), and bFGF (PeproTech, 100-18B) supplemented at a concentration of 20 ng/mL. The plate was left at 37 °C for 3 hours after which the ROCK-I containing seeding media was removed. The cells were then transfected as described above using NutriStem (100 µL/well) without ROCK-I. The cells were incubated with DNA-Transfection reagent mixture for 24 h until the micropatterns reached confluency. For the experiment where we tested exogenous BMP4, medium was replaced with NutriStem and left in Nutristem for 24 h until they reached confluency. At this point differentiation was induced by replacing NutriStem with the N2B27 media supplemented with either BMP4 and NODAL or only NODAL (for transfected samples). NODAL (R&D Systems, Catalog # 3218-ND) was added at 100 ng/mL. BMP4 (R&D Systems, Catalog # 314-BP-MTO) was used at a concentration of 50 ng/mL. For negative control, BMP4 was not added to the N2B27 media. The volume of media used for differentiation was 150 µL/well. 36 h after induction of differentiation, the cell culture supernatant was collected for ELISA, 3 replicas were harvested for live flow cytometry and remaining replicas (12 for each condition) were fixed and subsequently stained with DAPI and TBXT and measured by confocal microscopy.

### Confocal imaging of micropatterned colonies

Micropatterned samples were fixed with 4 % paraformaldehyde and permeabilized using 100 % cold methanol. Cells were exposed to the Human/Mouse Brachyury Affinity Purified Polyclonal Goat IgG antibody (R&D Systems, Catalog # AF2085) overnight at 4 °C. The antibody was incubated in 2% FBS in HBSS. The samples were washed and immersed in the secondary antibody solution containing DAPI for 90 minutes at room temperature and subsequently rinsed.

Images were acquired using the Zeiss LSM 800 Confocal Microscope. We used magnification of 20 x and obtained 5 z-slices/colony. The RUES2 cell line contains endogenous tags for the markers SOX2 (mCitrine), SOX17 (tdTomato), and TBXT (mCerulean). We imaged SOX2 using the 488 nm diode laser (545 short pass filter) and SOX17 via the 561 nm diode laser (610 short-pass filter). Due to the mCerulean signal being faint, we stained TBXT with the AlexaFluor 647 secondary antibody and imaged the signal with the 647 nm diode laser (655 long-pass filter). Finally, the 647 nm diode laser was used to image the iRFP tag.

### ELISA

BMP4 concentration released into the media was quantified using the BMP-4 Human ELISA Kit from (ThermoFisher, EHBMP4). Standards and protocols were done according to the instructions of the kit. The ELISA results were analyzed for absorbance at both 450nm and 550 nm using the Tecan Infinite M Plex and standard curve was calculated using CurveExpert software.

### Flow Cytometry

Cells were incubated with TrypLE for 5 minutes at 37 °C, mixed with equal volume of growth medium, dissociated to single cells by pipetting and transferred to 96-well v-bottom plate for dead/live and / or antibody staining. Cells were spun down at 1200rpm (335G, Rotor Sx4750, Beckman Coulter Allegra X-12R centrifuge) for 3 minutes and washed once with HF Buffer (HBSS, Gibco, 14175103) supplemented with 4 % FBS. Cells that were not stained with antibodies, (which were all transfections in 96-well plates in pluripotency state) were incubated with a dead / live stain chosen based on compatibility with fluorescent reporter combination. Dead/live stains used were DAPI (Sigma-Aldrich, D9542) incubated at a concentration of 0.1 µg/mL in HF for 3 minutes at room temperature or Zombie NIR^TM^ (BioLegend, 423106) incubated in PBS at a 1:500 dilution for 15 minutes at room temperature. Cells have then been washed once with HF, suspended to single cells and used for Flow Cytometry.

For definitive endoderm staining, half of a well from a 12-well plate (2-4 x10^5^ cells) were dissociated, washed once with HF and co-stained with conjugated PE-CY7 anti-human CD184 (BD, 560669) and APC anti-human CD117 (BD, 550412) or with the corresponding isotype controls PE-CY7 Mouse Ig2akappa Isotype (BD, 557907) and APC mouse IgG1kappa Isotype (BD, 555751). Each antibody was diluted to 1:200 and incubated in PBS containing LIVE/DEAD™ Fixable Violet Dead Cell Stain (diluted 1:1000) for 30 minutes at room temperature. Cells were washed twice with HF buffer before using for Flow Cytometry.

For pluripotency markers staining, all cells from a 12-well (4-6×10^5^ cells) were harvested and stained with LIVE/DEAD™ Fixable Violet Dead Cell Stain (diluted 1:1000) for 30 minutes at room temperature. Cells were washed in HF buffer and incubated in 4 % Formaldehyde diluted in HF Buffer for 15 minutes at room temperature. Cells were washed with HF and stored up to one week at 4 °C if necessary. For permeabilization, cells were gently resuspended in 100% ice-cold Methanol. After 2 minutes, twice the volume of HF was added to the cells and spun down at 1200 rpm for 3 minutes. Then cells were resuspended with primary antibodies anti-Oct-3/4 (BD, 611203) and anti-Sox2 (D6D9) (Cell Signaling Technologies, 3579S) diluted 1:100 in HF buffer and incubated for 30 minutes at room temperature. Cell were washed three times with HF buffer and incubated with secondary antibodies APC-Cy7 goat anti-rabbit IgG (BD, 611203) used at 1:200 dilution and AF647 Donkey anti-mouse IgG (Thermo Fisher Scientific, A-31571) used at 1:400 dilution and incubated for 15 minutes at room temperature. Negative background controls with only secondary antibodies were generated. Cells were washed once with HF and used for Flow Cytometry.

For both, pluripotency and endoderm antibody staining, single antibody-stained controls, unstained controls with and without Dead / Live stain and single Dead / Live stain controls using ArC™ Amine Reactive Compensation Beads (Thermo Fisher Scientific, A10628) were prepared for gating and cross-talk evaluation.

Flow Cytometry was performed using BD LSR Fortessa Analyzer for samples shown in Figure 2 to Figure 6. Micropatterned RUES2 cells (Figure 7) were measured using Beckman Coulter CytoFLEX LX flow cytometer. The laser-filter configuration and PMT voltage used for each fluorophore can be found in Table S6. We used Calibration beads (BD, 642412) to ensure consistent machine performance and SPHERO RainBow Calibration particles (BD, 559123) to track and correct for consistency of PMT settings for each channel across different experiments.

### Analysis of Flow Cytometry data

All flow cytometry data were analyzed using FlowJo software. Single color controls were used to estimate the crosstalk of each fluorophore into each channel. Compensation was performed using the FlowJo compensation matrix if necessary. The values in the various charts, labelled as relative expression units (rel.U), were calculated as follows: (I) First single and live cells were gated based on their forward and side scatter readouts and the absence of dead/live marker. (ii) From this selection, cells that are positive in a given fluorophore were gated using fluorophore-negative, transfection marker-positive single-color control such that 99.9% of cells in this single-color control sample fall outside of the selected gate. (iii) For each positive cell population in a given channel, the mean value of the fluorescent intensity was calculated and multiplied by the frequency of the positive cells. This value was used as a measure for the total reporter signal in a sample. The total reporter signal of a circuit output was then normalized with the total signal of the reporter that has been used as a transfection control to counterbalance possible transfection variation. The procedure can be summed in the following formula:

Reporter intensity of a sample in relative units (rel.U.) = [mean (Reporter in Reporter+ cells) × Frequency (Reporter+ cells)]/[mean (Transfection Marker in Transfection Marker+ cells) × frequency (Transfection Marker+ cells)].

Reporter intensity in absolute unit (a.u.) as in Figure 7B and 7E = [mean (iRFP Reporter in Reporter+ cells) × Frequency (iRFP Reporter+ cells)]

Frequency of cells expressing a given marker as in Figure S2C and S2D, were calculated from I) single and live cells were gated based on their forward and side scatter readouts and the absence of dead/live marker. (ii) From this selection, cells that are positive in a given marker were gated using control samples generated either with samples stained with only secondary antibodies (Figure S2C) and isotype controls (Figure S2D), respectively.

Data points in figures are shown as mean ± standard deviation (s.d.) of three or more independent biological samples. A two-sided unpaired *t*-test was performed on selected sample combinations as indicated in figures. Homoscedastic or heteroscedastic t-tests were performed based on the results of an F-test using a P-value of 0.05. *P* values of the t-test are indicated in figures as required.

Viability in Figure S2B is calculated from the fraction of “live” cells in the Side and Forward Scatter multiplied with the fraction of cells that are negative for LIVE/DEAD^TM^ Violet stain. Correlation plots in Figure 5D were created with a simple linear regression model of the form y = a + b*x. The regression coefficient (R^2^) was calculated to evaluate which circuit fits better with model predictions.

### Analysis of micropatterned colonies

CellProfiler was used to identify nuclear regions based on the intensities in the DAPI images. The intensities of Sox2, Sox17 and TBXT were then measured within these nuclear regions for each channel. Next, the single-cell data of nuclear location and protein intensity was analysed using ContextExplorer (CE) [65] as described elsewhere [73]. Shortly, using the DBSCAN algorithm in CE we clustered cells into colonies and assigned xy-coordinates to each cell relative to the colony center. Cells were then grouped into hexagonal or annular bins to create the aggregation or line plots, respectively, and calculated the mean intensities and standard error of the markers within these bins. To analyze the differences in germ-layer marker expression (e.g. whether more cells exhibited SOX17 for a certain tuner or condition vs others), we calculated ‘area under the curve’ (AUC) for the radial profiles of each marker. We first normalized the intensity values calculated via CE such that the highest radial intensity = 1. Next, we calculated the ‘area under the curve’ (and the associated standard error) using the lowest radial intensity as baseline. Therefore, each condition/tuner had its unique baseline dependent on the marker expression. The ‘area under the curve’ analysis were performed in GraphPad Prism 8.1. Table S6 summarized the mumber of colonies analyzed for each sample.

## Materials, data and code availability

Plasmids generated in this study are in process of being submitted to Addgene as indicated in Table S5. The code for automated miRNA identification with modified pre-processing pipeline is available on GitHub: https://github.com/jwon0408/SynNetModified. Unprocessed data are summarized in Table S2, S3 and S6. Additional raw data can be shared upon request.

## Author contributions

Conceptualization: L.P., Y.S.M, P.W.Z.; Methodology: L.P.; Vector Design: L.P, C.L.; Vector cloning: L.P., C.L., D.W., E.Y., M.V.; Circuit design: L.P.; miRNA identification:, E.Y., Y.S.M, R.D.J, L.P, Circuit Modeling: D.W., L.P.; Cell culture: L.P., T.Y., H.K.; Differentiations: L.P., T.Y.; Transfections: L.P., T.Y., Y.S.M, D.W., M.V.; Flow Cytometry L.P., Y.S.M., C.L., T.Y.; Micropatterning: H.K.; Confocal Microscopy: M.S., H.K.; ELISA: C.L.; Data curation: L.P.; Data analysis: L.P., C.L., H.K., Resources: P.W.Z., Y.B., P.G.; Writing - original draft: L.P.; Writing - review & editing: L.P., Y.S.M., P.G., P.W.Z., H.K., M.V., D.W., E.Y., Y.B., R.D.J; Visualization: L.P.; Project administration: L.P., T.Y., P.W.Z.; Funding acquisition: L.P., P.W.Z, H.K., Y.B.

## Acknowledgements

The authors would like to thank Dr. Geoffrey Clarke for discussion on computational modeling, Prof. Charles King for discussion and providing raw miRNA Sequencing data. Prof. Derek Van der Kooy and Prof. Penney Gilbert for sharing lab space and infrastructure. Prof. Christina Nostro for providing hESC H1 cell line. Prof. Martin Fussenegger for providing the plasmids encoding PIT2 and ET1 activators and their regulated promoter sequences. We also want to thank Dionne White from Temetry Faculty of Medicine Flow Cytometry Facility at U of T for troubleshooting technical difficulties with Flow Cytometry. Lastly, we would like to thank all Zandstra lab members for discussions.

