## Supplementary Information for "Discrete-to-analog signal conversion in human pluripotent stem cells"

Figures S1-S7

Tables S4 and S5
(Tables S1,S2,S3 and S6 in separate excel files)

### Figure S1

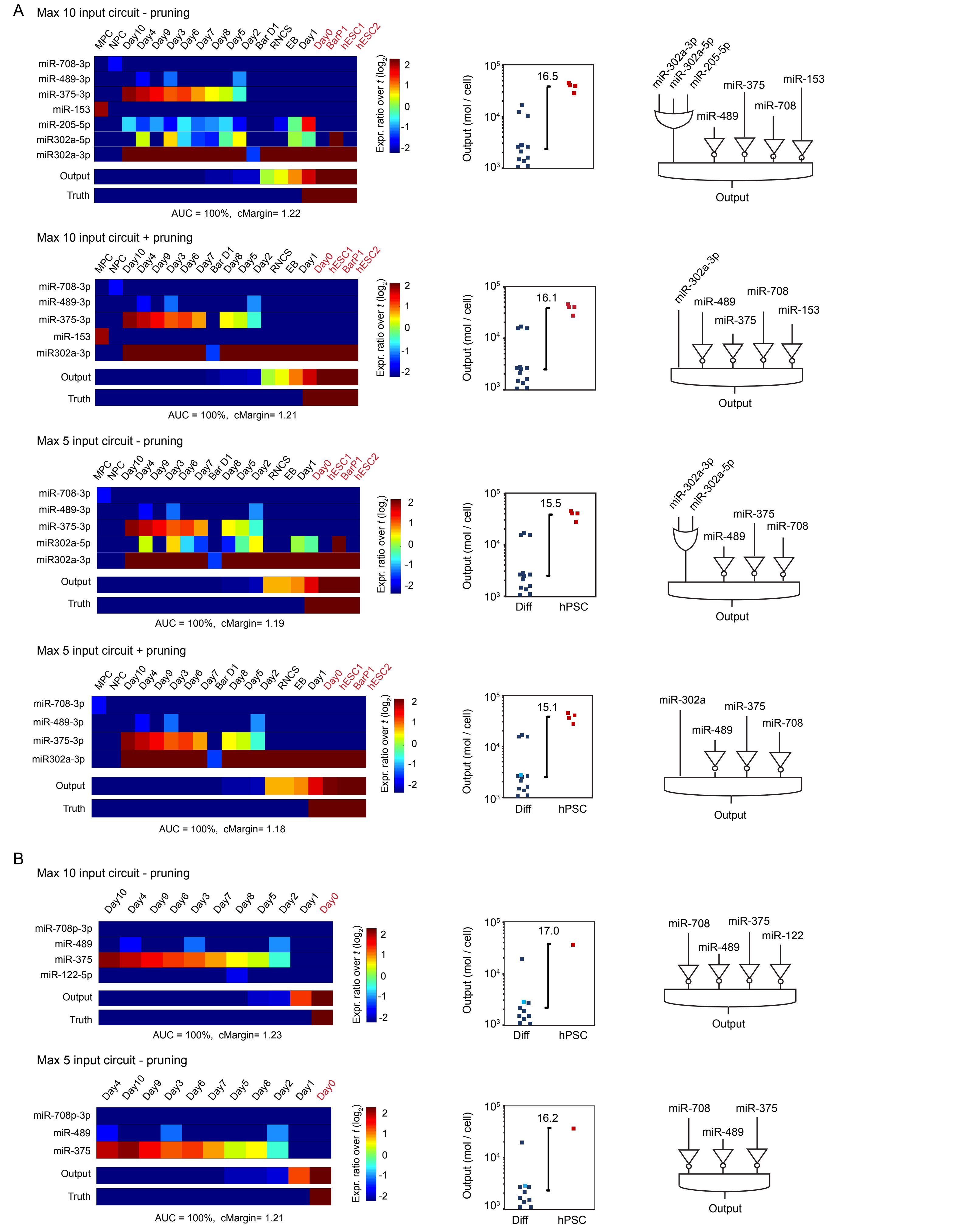

A
Max 10 input circuit - pruning
MPC NPC Day10 Day4 Day9 Day3 Day6 Day7 Day8 Day5 Day2 Bar D1 RNCS EB Day1 Day0 BarP1 hESC1 hESC2
miR-708-3p
miR-489-3p
miR-375-3p
miR-153
miR-205-5p
miR302a-5p
miR302a-3p
Output
Truth
AUC = 100%, cMargin= 1.22
Expr. ratio over t (log2)
2 1 0 -1 -2
Output (mol / cell)
10^5 10^4 10^3
16.5
miR-302a-3p
miR-302a-5p
miR-205-5p
miR-489
miR-375
miR-708
miR-153
Output
Max 10 input circuit + pruning
MPC NPC Day10 Day4 Day9 Day3 Day6 Day7 Bar D1 Day8 Day5 Day2 RNCS EB Day1 Day0 hESC1 BarP1 hESC2
miR-708-3p
miR-489-3p
miR-375-3p
miR-153
miR302a-3p
Output
Truth
AUC = 100%, cMargin= 1.21
16.1
miR-302a-3p
miR-489
miR-375
miR-708
miR-153
Output
Max 5 input circuit - pruning
MPC NPC Day10 Day4 Day9 Day3 Day6 Day7 Bar D1 Day8 Day5 Day2 RNCS EB Day1 Day0 hESC1 BarP1 hESC2
miR-708-3p
miR-489-3p
miR-375-3p
miR302a-5p
miR302a-3p
Output
Truth
AUC = 100%, cMargin= 1.19
15.5
Diff
hPSC
miR-302a-3p
miR-302a-5p
miR-489
miR-375
miR-708
Output
Max 5 input circuit + pruning
MPC NPC Day10 Day4 Day9 Day3 Day6 Day7 Bar D1 Day8 Day5 Day2 RNCS EB Day1 Day0 hESC1 BarP1 hESC2
miR-708-3p
miR-489-3p
miR-375-3p
miR302a-3p
Output
Truth
AUC = 100%, cMargin= 1.18
15.1
Diff
hPSC
miR-302a
miR-489
miR-375
miR-708
Output
B
Max 10 input circuit - pruning
Day10 Day4 Day9 Day6 Day3 Day7 Day8 Day5 Day2 Day1 Day0
miR-708p-3p
miR-489
miR-375
miR-122-5p
Output
Truth
AUC = 100%, cMargin= 1.23
17.0
Diff
hPSC
miR-708
miR-489
miR-375
miR-122
Output
Max 5 input circuit - pruning
Day4 Day10 Day9 Day3 Day6 Day7 Day5 Day8 Day2 Day1 Day0
miR-708p-3p
miR-489
miR-375
Output
Truth
AUC = 100%, cMargin= 1.21
16.2
Diff
hPSC
miR-708
miR-489
miR-375
Output

**Figure S1** Summary of computationally identified circuits showing input miRNAs and predicted circuit output levels using miRNA data sets from Fogel et al, Lipchina et al and Bar et al. **(A**) or only Fogel et al **(B)**. For both A and B, expression levels of identified miRNAs inputs are given as fold change over the pre-set input abundance threshold (t) of the total miRNA pool (where t = 1%) (left). Calculated circuit output levels are given as mol / cell (middle) and logic connectivity of the identified miRNA is depicted (right). miRNA expression data and nomenclature can be found in TableS1, raw output data and constraint files of the algorithm in Table S2 for A and Table S3 for B. Related to Figure 2.

Figure S2

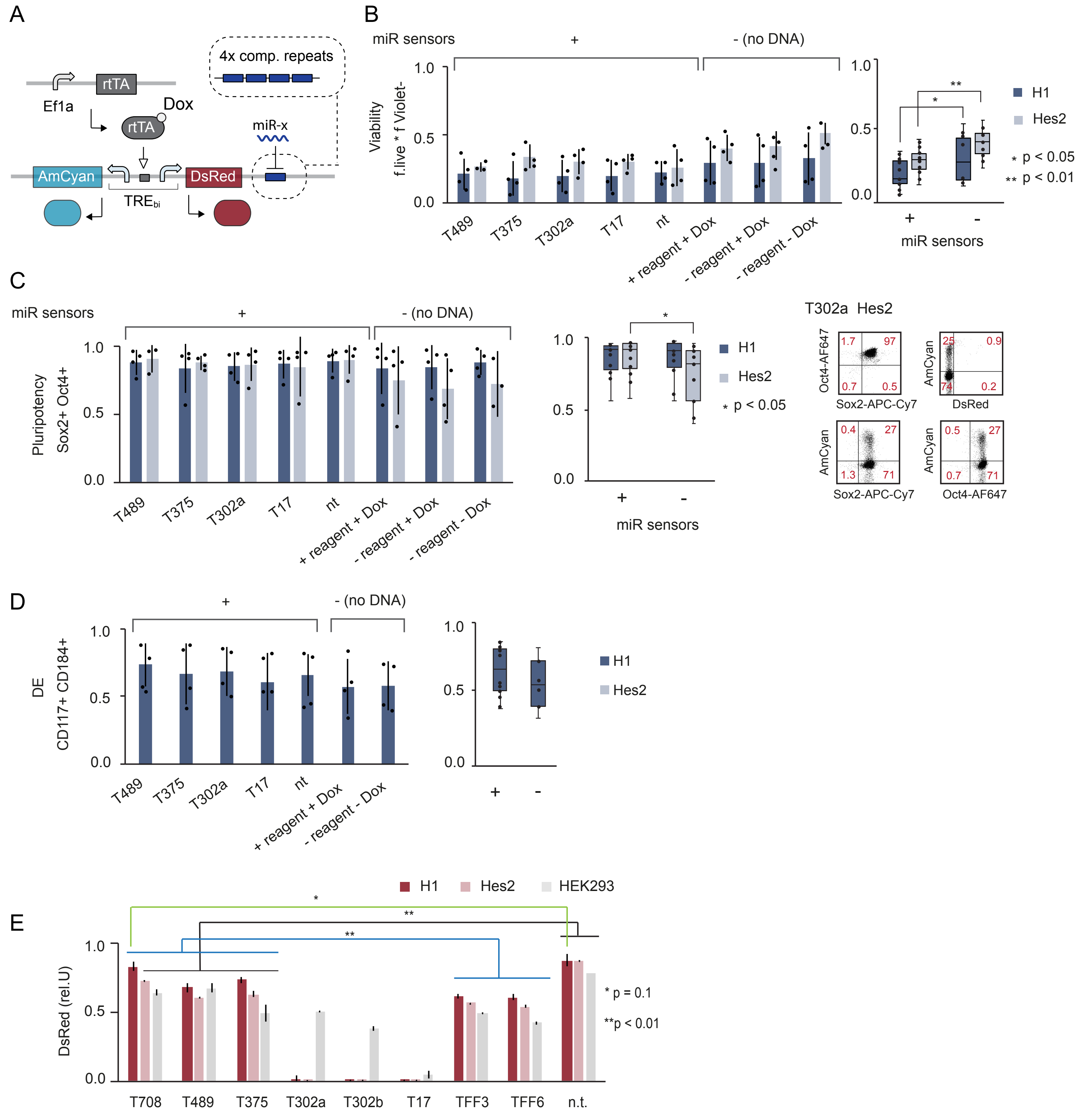
A
4x comp. repeats
rtTA
Ef1a
Dox
rtTA
miR-x
AmCyan
DsRed
TREbi
B
miR sensors
+
- (no DNA)
Viability
f.live * f Violet-
1.0
0.5
0.0
T489
T375
T302a
T17
nt
+ reagent + Dox
- reagent + Dox
- reagent - Dox
**
*
H1
Hes2
* p < 0.05
** p < 0.01
+
-
miR sensors
C
miR sensors
+
- (no DNA)
Pluripotency
Sox2+ Oct4+
1.0
0.5
0.0
T489
T375
T302a
T17
nt
+ reagent + Dox
- reagent + Dox
- reagent - Dox
*
H1
Hes2
* p < 0.05
+
-
miR sensors
T302a Hes2
Oct4-AF647
1.7
97
0.7
0.5
Sox2-APC-Cy7
AmCyan
25
0.9
74
0.2
DsRed
AmCyan
0.4
27
1.3
71
Sox2-APC-Cy7
AmCyan
0.5
27
0.7
71
Oct4-AF647
D
+
- (no DNA)
DE
CD117+ CD184+
1.0
0.5
0.0
T489
T375
T302a
T17
nt
+ reagent + Dox
- reagent - Dox
H1
Hes2
+
-
E
H1
Hes2
HEK293
*
**
**
DsRed (rel.U)
1.0
0.5
0.0
T708
T489
T375
T302a
T302b
T17
TFF3
TFF6
n.t.
* p = 0.1
**p < 0.01

**Figure S2 A)** Illustration of bi-directional miRNA sensor system. **B - D)** Bar chart left showing the fraction of Oc4+ Sox+ double positive pluripotent cells (B), viability (C) and CD184+ and CD117+ double positive cells marking definitive endoderm (DE) (D) for each miRNA sensor (+) and the untransfected samples (-) in different conditions as indicated. Individual samples are indicated as black circles. Bar chart right shows all samples transfected with miRNA sensors (+) with all untransfected samples (-) in H1 and HES-2. Median (horizontal line in box) and s.d.(error bars) are indicated. **E)** Bar chart showing relative DsRed expression of sensors containing the indicated miRNA target sites as four fully complementary repeats . The control vector (n.t.) does not contain any target sites. Two-tailed unpaired t-tests were performed to compare T708, T489 and T375 to TFF3,TFF6 or to n.t.. for each cell line. P values are shown as indicated. The bar groups samples with identical P-values. Each bar chart corresponds to mean +/- s.d. from at least three biological replicates. Related to Figure 3.

### Figure S3

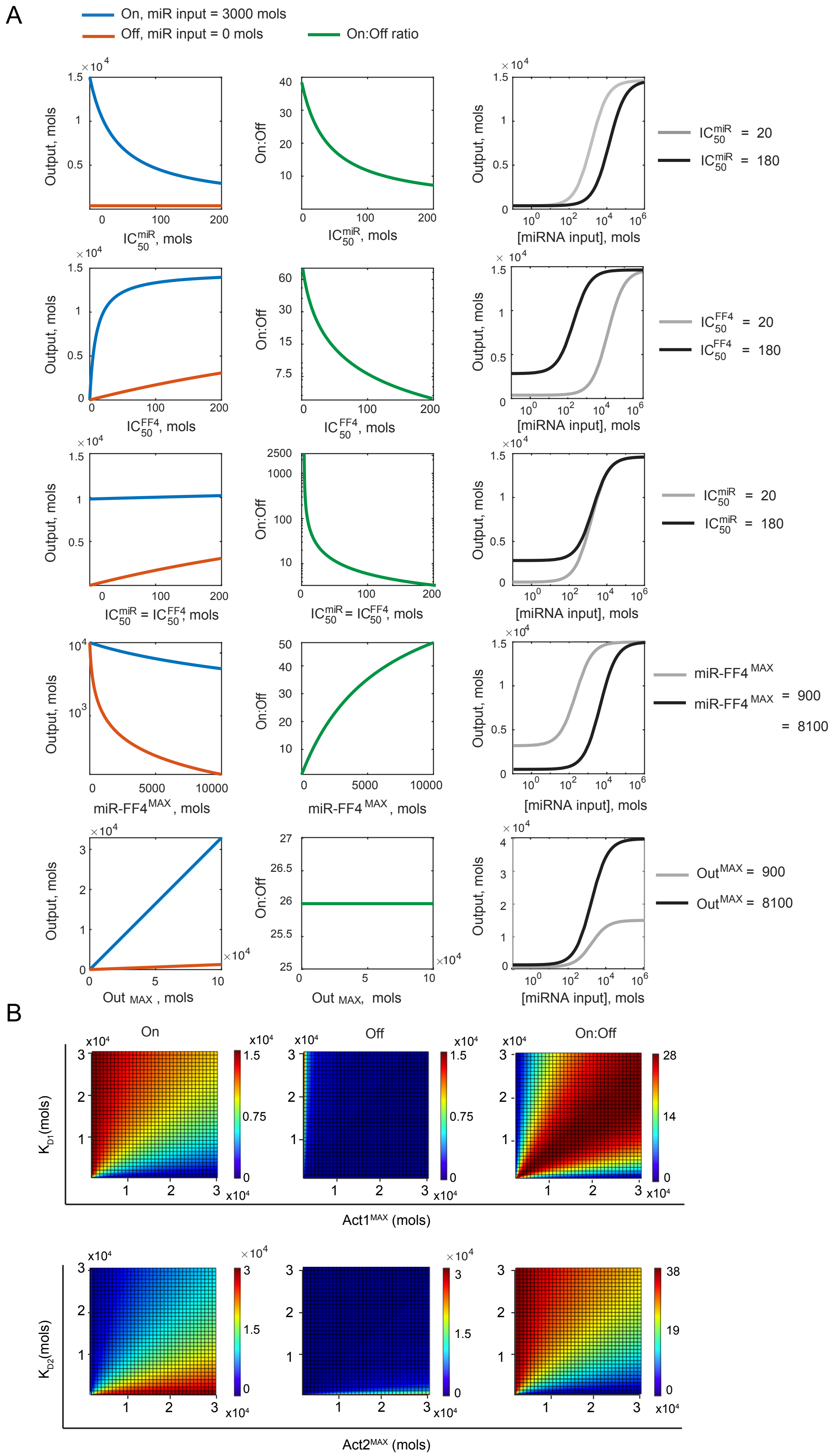

A
On, miR input = 3000 mols
Off, miR input = 0 mols
On:Off ratio
Output, mols
On:Off
IC50 miR, mols
[miRNA input], mols
IC50 miR = 20
IC50 miR = 180
IC50 FF4, mols
IC50 FF4 = 20
IC50 FF4 = 180
IC50 miR = IC50 FF4, mols
miR-FF4 MAX, mols
miR-FF4 MAX = 900
miR-FF4 MAX = 8100
Out MAX, mols
Out MAX = 900
Out MAX = 8100
B
On
Off
On:Off
KD1(mols)
Act1MAX (mols)
KD2(mols)
Act2MAX (mols)

**Figure S3 A)** Single parameter screening. Shown are dose response curves of individual parameters ($IC_{50}^{miR}$, $IC_{50}^{FF4,}$ miR-FF4$^{MAX}$ and Out$^{MAX}$) for On and Off state (left), On / Off ratio (middle) and input – output transfer function for each parameter at low and high miRNA levels (right). **B)** Combinatorial screening of KD and Act$^{MAX}$ for Act1 (top) and Act2 (bottom). Shown are calculated output levels in On state (left), Off state (middle) and On / Off ratio. Related to Figure 4.

Figure S4

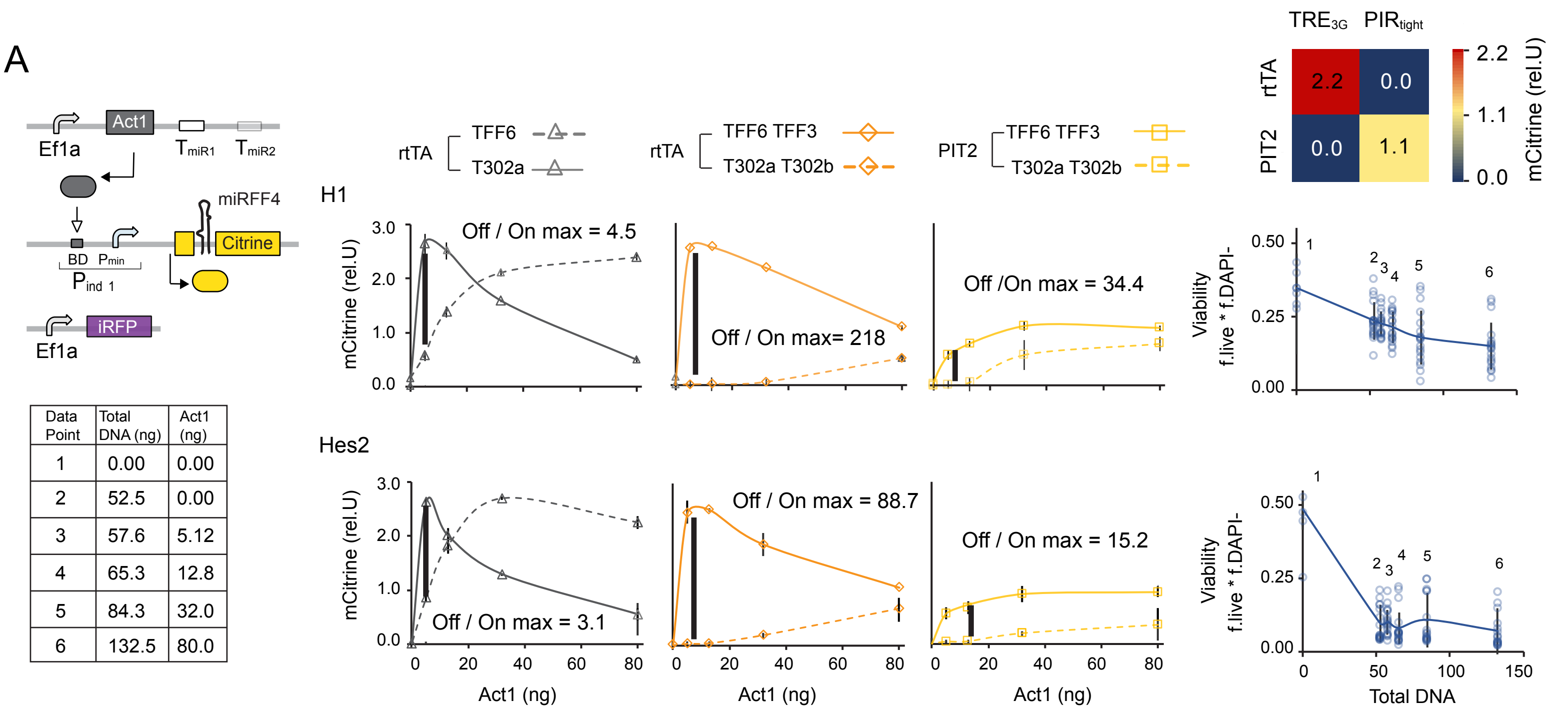


| Data Point | Total DNA (ng) | Act1 (ng) |
|---|---|---|
| 1 | 0.00 | 0.00 |
| 2 | 52.5 | 0.00 |
| 3 | 57.6 | 5.12 |
| 4 | 65.3 | 12.8 |
| 5 | 84.3 | 32.0 |
| 6 | 132.5 | 80.0 |

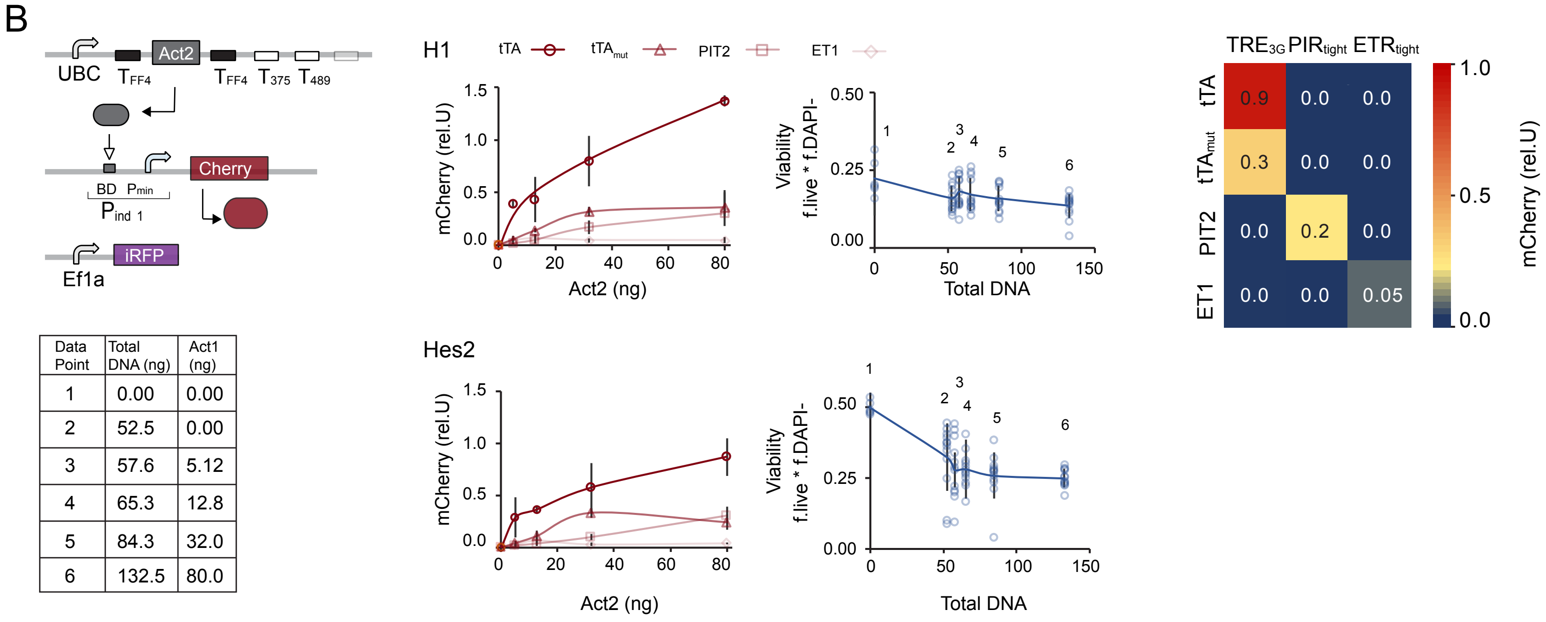


| Data Point | Total DNA (ng) | Act1 (ng) |
|---|---|---|
| 1 | 0.00 | 0.00 |
| 2 | 52.5 | 0.00 |
| 3 | 57.6 | 5.12 |
| 4 | 65.3 | 12.8 |
| 5 | 84.3 | 32.0 |
| 6 | 132.5 | 80.0 |

**Figure S4** Characterization of the FF4 inducible **(A)** and output inducible **(B)** module, respectively. Dose response function (middle) showing normalized fluorescence readouts of mCitrine and mCherry to the changing Act1 and Act2 plasmid amounts, respectively. Curves serve as visual guides. Charts show the mean ± s.d. of at least three biological replicates. Viability as a function of DNA amount (left). Single data from all samples are plotted as circles. Mean values are indicated as dots. Line serves as a visual guide connecting mean values. Color plots show cross-activation of each transactivator with each promoter using 32 ng of a given transactivator. Experimental description and raw data are provided in Table S6. Related to Figure 5.

### Figure S5

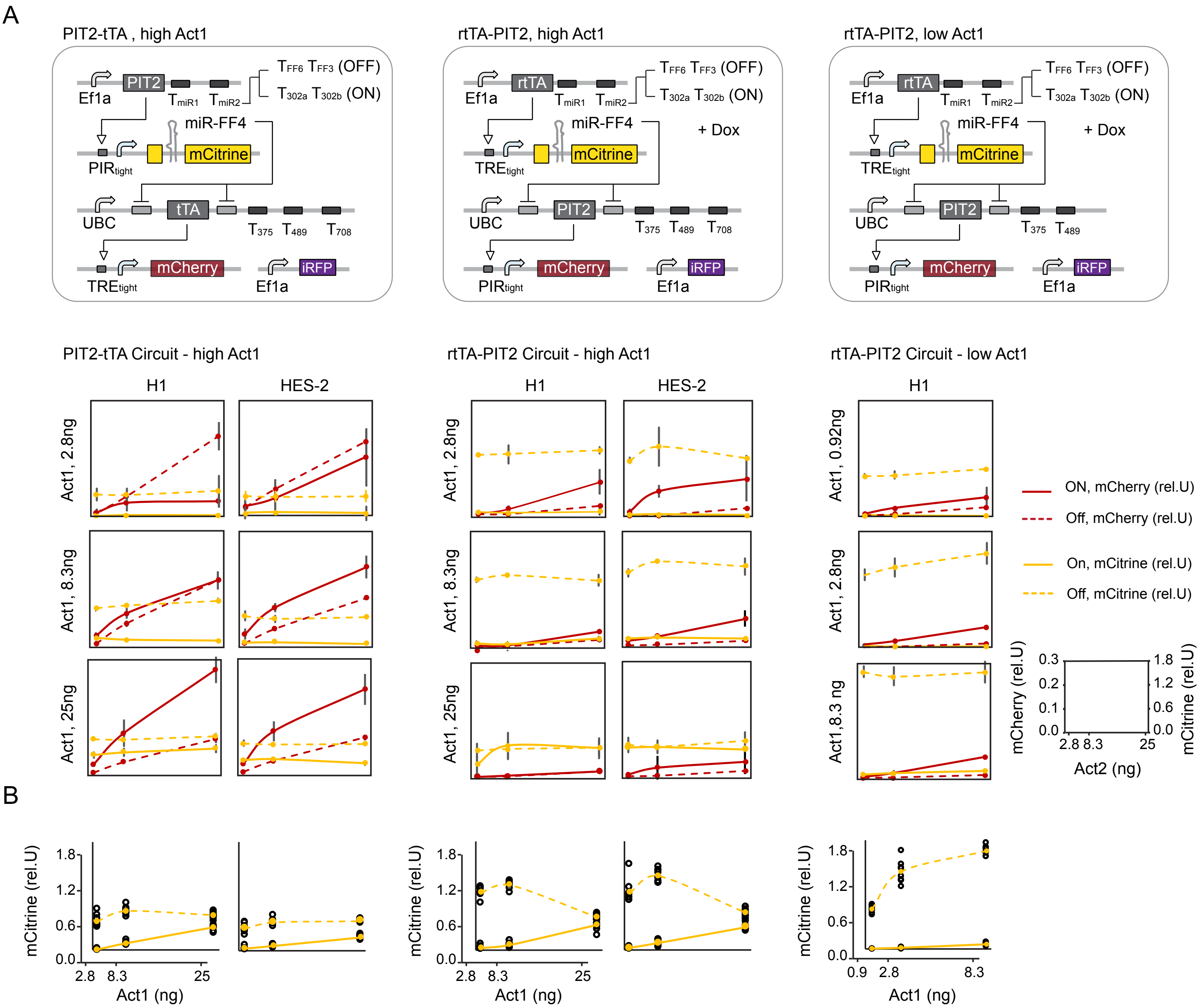

A
PIT2-tTA , high Act1
rtTA-PIT2, high Act1
rtTA-PIT2, low Act1
Ef1a
PIT2
rtTA
miR-FF4
mCitrine
tTA
UBC
mCherry
iRFP
+ Dox
PIT2-tTA Circuit - high Act1
rtTA-PIT2 Circuit - high Act1
rtTA-PIT2 Circuit - low Act1
H1
HES-2
Act1, 2.8ng
Act1, 8.3ng
Act1, 25ng
Act1, 0.92ng
Act1,8.3 ng
ON, mCherry (rel.U)
Off, mCherry (rel.U)
On, mCitrine (rel.U)
Off, mCitrine (rel.U)
mCherry (rel.U)
mCitrine (rel.U)
Act2 (ng)
B
mCitrine (rel.U)
Act1 (ng)

**Figure S5 A)** Experimental combinatorial screening of Act1 and Act2 in three different circuit configurations: PIT2-tTA using high levels of Act1 (left, data shown in Figure 5C left), rtTA-PIT2 using high levels of Act1 (middle) and rtTA-PIT2 using low levels of Act1 (right, data shown in Figure 5C right). For each circuit and each Act1 concentration, the normalized mCitrine / FF4 level (yellow) and the normalized Output levels (mCherry) (red) are shown in response to changing amount of Act2 expressing plasmids in the On configuration (Act1 targeted by TFF3/TFF6) (solid lines) and Off configuration (Act1 targeted by T302a/T302b) (dashed lines). **B)** Scatter plot showing normalized mCitrine expression in response to changing amounts of Act1 expressing plasmids across all Act2 levels. Line connects mean values and serves as a visual guide. All charts show mean ± s.d. of at least three biological replicates. Experimental description and raw data are provided in Table S6. Related to Figure 5A-D.

Figure S6

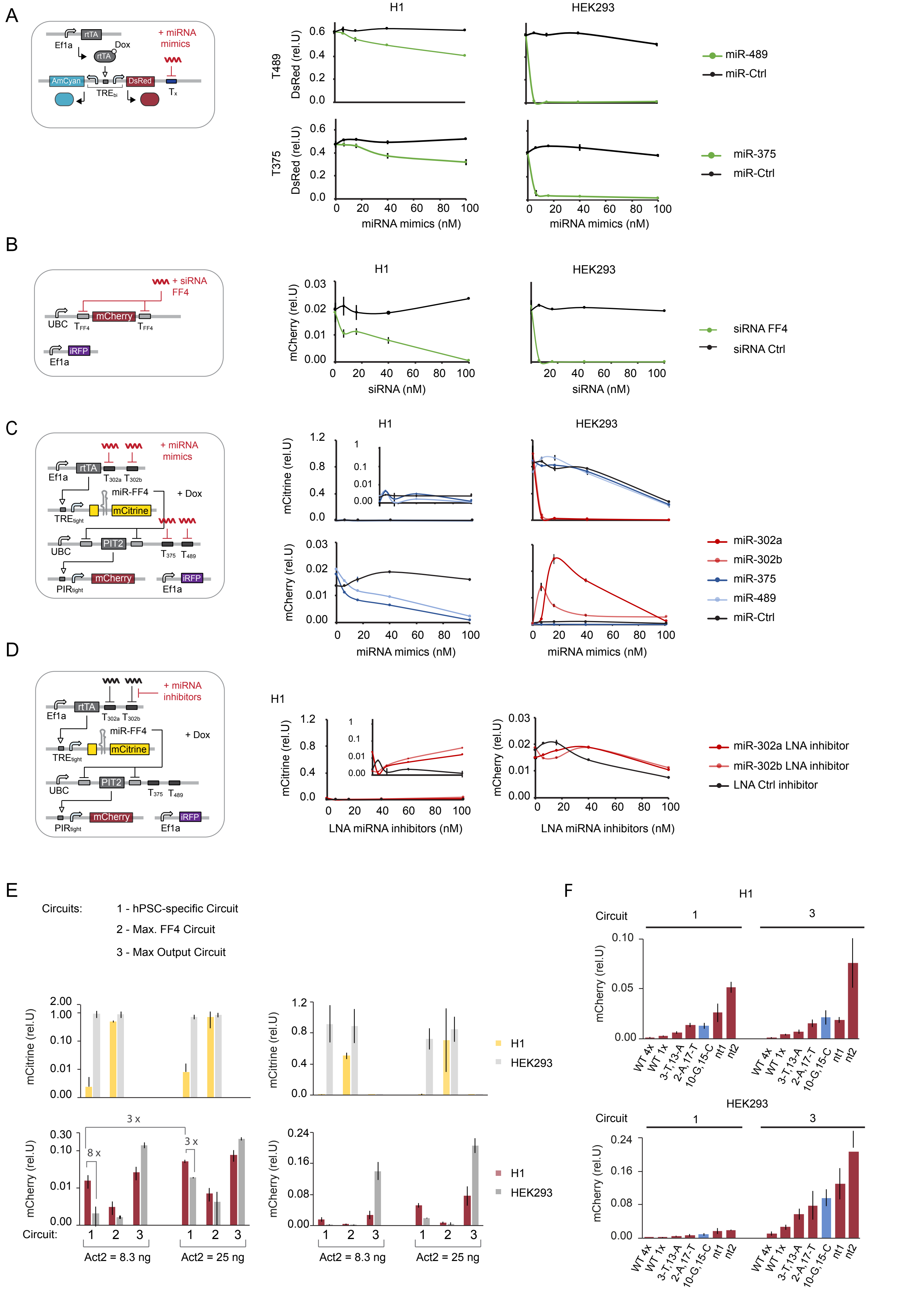

A
Ef1a
rtTA
Dox
+ miRNA mimics
AmCyan
TREbi
DsRed
Tx
H1
HEK293
T489
DsRed (rel.U)
miR-489
miR-Ctrl
T375
miR-375
miRNA mimics (nM)
B
+ siRNA FF4
UBC
TFF4
mCherry
iRFP
mCherry (rel.U)
siRNA FF4
siRNA Ctrl
siRNA (nM)
C
T302a T302b
miR-FF4
+ Dox
TREtight
mCitrine
PIT2
T375 T489
PIRtight
mCitrine (rel.U)
miR-302a
miR-302b
miR-375
miR-489
miR-Ctrl
D
+ miRNA inhibitors
miR-302a LNA inhibitor
miR-302b LNA inhibitor
LNA Ctrl inhibitor
LNA miRNA inhibitors (nM)
E
Circuits:
1 - hPSC-specific Circuit
2 - Max. FF4 Circuit
3 - Max Output Circuit
H1
HEK293
3 x
8 x
3 x
Circuit:
Act2 = 8.3 ng
Act2 = 25 ng
F
Circuit
WT 4x
WT 1x
3-T,13-A
2-A,17-T
10-G,15-C
nt1
nt2

**Figure S6 A)** Repression of bi-directional reporter by miR-375 and miR-489 mimics. DsRed is either targeted with four fully complementary miR-489 (top) or miR-375 (bottom) targets sites. Charts show normalized DsRed expression in response to increasing amounts of miRNA mimics in H1 and HEK293 as indicated. **B)** Repression strength of siRNA FF4 on UBC driven mCherry reporter flanked with FF4 targets sites (same design as Act2 expressing constructs in Figure 5 and 6). Chart shows normalized mCherry expression in response to increasing amounts of siRNA FF4 **C)** hPSC-specific circuit (circuit 1) in response to miRNA mimic administration. Charts show mCitrine (top) and mCherry (bottom) expression in response to increasing concentrations of miRNA mimics in H1 (left) and HEK293 (right). Related to Figure 6D. **D)** hPSC-specific circuit (circuit 1) in response to LNA inhibitors. Charts show mCitrine (top) and mCherry (bottom) expression in response to increasing amounts of miR-302a and miR-302b LNA inhibitors in H1 (left) and HEK293 (right). **E)** Comparison of circuit performance using 8.3 and 25 ng of Act2-expressing plasmid, corresponding to a 0.1:1:0.3:1 and a 0.1:1:1:1 molar ratio between circuit plasmids, respectively. Bar charts show FF4 / mCitrine expression (top) and output / mCherry expression (bottom) in log scale (left) and linear scale (right). Related to Figure 6C. **F)** Bar chart showing fine tuning of output using miSFITs library in H1 and HEK293. Related to Figure 6E. All charts show mean ± s.d. of at least three biological replicates. Related to Figure 6.

Figure S7

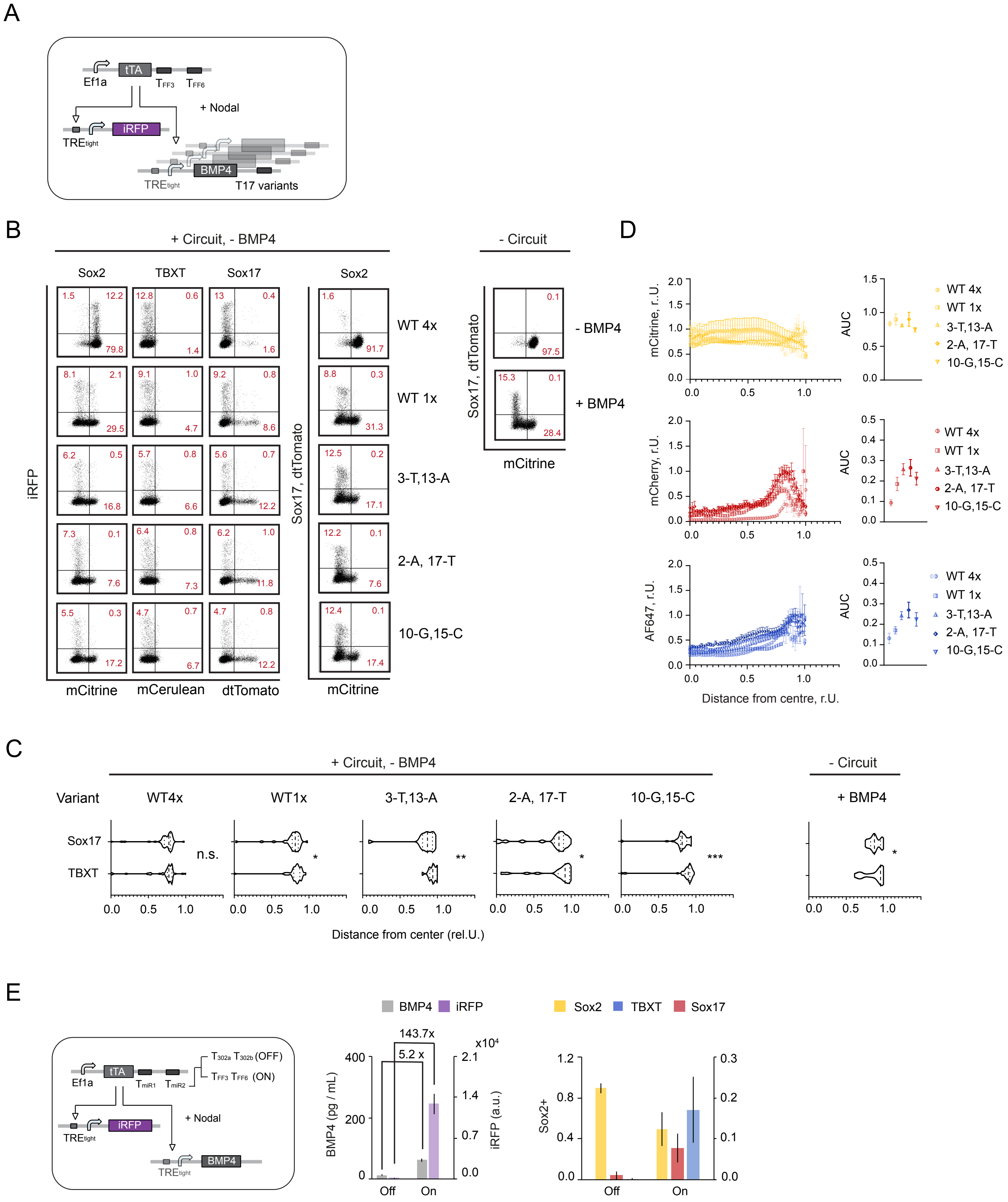

A
Ef1a
tTA
TFF3
TFF6
+ Nodal
iRFP
TREtight
BMP4
TREtight
T17 variants
B
+ Circuit, - BMP4
- Circuit
Sox2
TBXT
Sox17
Sox2
iRFP
Sox17, dtTomato
WT 4x
WT 1x
3-T,13-A
2-A, 17-T
10-G,15-C
- BMP4
+ BMP4
mCitrine
mCerulean
dtTomato
mCitrine
mCitrine
C
+ Circuit, - BMP4
- Circuit
Variant
WT4x
WT1x
3-T,13-A
2-A, 17-T
10-G,15-C
+ BMP4
Sox17
TBXT
n.s.
Distance from center (rel.U.)
D
mCitrine, r.U.
mCherry, r.U.
AF647, r.U.
AUC
WT 4x
WT 1x
3-T,13-A
2-A, 17-T
10-G,15-C
Distance from centre, r.U.
E
Ef1a
tTA
TmiR1
TmiR2
T302a T302b (OFF)
TFF3 TFF6 (ON)
+ Nodal
iRFP
TREtight
BMP4
TREtight
BMP4
iRFP
143.7x
5.2 x
x10⁴
BMP4 (pg / mL)
iRFP (a.u.)
Off
On
Sox2
TBXT
Sox17
Sox2+
Off
On

**Figure S7** **A)** Circuit schematic. **B)** Representative Flow Cytometry scatter plots for microscopy data depicted in Figure 7C. **C)** Spatial divergence of Sox17 and TBXT. **D)** Radial marker profiles of Sox2, TBXT and Sox17 for all miSFITs variants (left) and associated AUC values (right). Charts show average ± standard error calculated from at least 20 colonies. **E)** Control of BMP4 in response to endogenous miRNAs. Micropatterned RUES2 have been transfected with illustrated circuits using tTA at 2.8 ng / 96 well (in contrast, data in Figure 7E used tTA construct at 25 ng / 96 well). Bar chart on the left shows average BMP4 concentration released from each circuit as determined by ELISA and absolute iRFP expression as measured by Flow Cytometry. Each bar corresponds to mean +/- s.d. from three biological replicates. Bar chart on the right shows the average fraction of cells expressing a given germ-layer marker analyzed from at least 20 colonies from Confocal Microscopy images. Related to Figure 7.

Table S4: miSFITs T17 variant target sequences

| Target _ID | Base change(s) | Target sequence 5' – 3' | Oligo sequence (incl. cloning overhangs) 5'-3' |
|---|---|---|---|
| V2 | 10-G, 15-C | ctacctgcaGtgtaCgcactttg | **ggccgcaaa**gaagaccaGCTTGCTACACGGGTCAACATACTAACACGCAActacctgc aGtgtaCgcactttgTCATCCTCCATACGCAGCTATGCggtgAGGGTActgtcttcagG |
| V4 | 2-A, 17-T | cAacctgcactgtaagTactttg | **ggccgcaaa**gaagaccaGCTTGCTACACGGGTCAACATACTAACACGCAAcAacctg cactgtaagTactttgTCATCCTCCATACGCAGCTATGCggtgAGGGTActgtcttcagG |
| V5 | 9-G | ctacctgcGctgtaagcactttg | **ggccgcaaa**gaagaccaGCTTGCTACACGGGTCAACATACTAACACGCAActacctgc GctgtaagcactttgTCATCCTCCATACGCAGCTATGCggtgAGGGTActgtcttcagG |
| V8 | 3-T; 13-A | ctTcctgcactgAaagcactttg | **ggccgcaaa**gaagaccaGCTTGCTACACGGGTCAACATACTAACACGCAActTcctgc actgAaagcactttgTCATCCTCCATACGCAGCTATGCggtgAGGGTActgtcttcagG |
| V11 | 9-G, 15-C | ctacctgcGctgtaCgcactttg | **ggccgcaaa**gaagaccaGCTTGCTACACGGGTCAACATACTAACACGCAActacctgc GctgtaCgcactttgTCATCCTCCATACGCAGCTATGCggtgAGGGTActgtcttcagG |
| V12 | 12-C, 21-G | ctacctgcactCtaagcactGtg | **ggccgcaaa**gaagaccaGCTTGCTACACGGGTCAACATACTAACACGCAActacctgc actCtaagcactGtgTCATCCTCCATACGCAGCTATGCggtgAGGGTActgtcttcagG |
| V15 | 3-C | ctCcctgcactgtaagcactttg | **ggccgcaaa**gaagaccaGCTTGCTACACGGGTCAACATACTAACACGCAActCcctgc actgtaagcactttgTCATCCTCCATACGCAGCTATGCggtgAGGGTActgtcttcagG |
| WT | None | Ctacctgcactgtaagcactttg | **gccgcaaa**gaagaccaGCTTGCTACACGGGTCAACATACTAACACGCAActacctgca ctgtaagcactttgTCATCCTCCATACGCAGCTATGCggtgAGGGTActgtcttcagG |

Table S5: Plasmid list

#### Bi-directional reporters

| Name | Description | Backbone | Source of insert | Deposit on Addgene |
|---|---|---|---|---|
| pLP110 | AmCyan-TRE-DsRed-T708-5p | pZ073 | ssDNA, Seq from miRBase | Yes |
| pLP111 | AmCyan-TRE-DsRed-T489-3p | pZ073 | ssDNA, Seq from miRBase | Yes |
| pLP112 | AmCyan-TRE-DsRed-T375-3p | pZ073 | ssDNA, Seq from miRBase | Yes |
| pLP113 | AmCyan-TRE-DsRed-T302a-3p | pZ073 | ssDNA, Seq from miRBase | Yes |
| pLP114 | AmCyan-TRE-DsRed-T302b-3p | pZ073 | ssDNA, Seq from miRBase | Yes |
| pLP115 | AmCyan-TRE-DsRed | pZ073 | Self-ligation | Yes |
| pLP110 | AmCyan-TRE-DsRed-T708-5p | pZ073 | ssDNA, Seq from miRBase | Yes |
| pLP111 | AmCyan-TRE-DsRed-T489-3p | pZ073 | ssDNA, Seq from miRBase | Yes |
| pLP118 | AmCyan-TRE-DsRed-TFF3 | pZ073 | ssDNA, Seq from Leisner et al.[1] | Yes |
| pLP119 | AmCyan-TRE-DsRed-TFF6 | pZ073 | ssDNA, Seq from Leisner et al.[1] | Yes |
| pLP176 | AmCyan-TRE-DsRed-T17 WT1x | pZ073 | ssDNA, Seq from miRBase | Yes |
| pLP177 | AmCyan-TRE-DsRed-T17 V4 | pZ073 | ssDNA, Seq from Michaels et al.[2] | Yes |
| pLP178 | AmCyan-TRE-DsRed-T17 V2 | pZ073 | ssDNA, Seq from Michaels et al.[2] | Yes |
| pLP179 | AmCyan-TRE-DsRed-T17 V5 | pZ073 | ssDNA, Seq from Michaels et al.[2] | Yes |
| pLP180 | AmCyan-TRE-DsRed-T17 V8 | pZ073 | ssDNA, Seq from Michaels et al.[2] | Yes |
| pLP181 | AmCyan-TRE-DsRed-T17 V7 | pZ073 | ssDNA, Seq from Michaels et al.[2] | Yes |
| pLP182 | AmCyan-TRE-DsRed-T17 V11 | pZ073 | ssDNA, Seq from Michaels et al.[2] | Yes |
| pLP183 | AmCyan-TRE-DsRed-T17 V12 | pZ073 | ssDNA, Seq from Michaels et al.[2] | Yes |
| pLP188 | AmCyan-TRE-DsRed-T17 WT | pZ073 | ssDNA, Seq from Michaels et al.[2] | Yes |
| pLP189 | AmCyan-TRE-DsRed-T17 V10 | pZ073 | ssDNA, Seq from Michaels et al.[2] | Yes |
| pLP190 | AmCyan-TRE-DsRed-V13 | pZ073 | ssDNA, Seq from Michaels et al.[2] | Yes |
| pLP191 | AmCyan-TRE-DsRed-neg Ctrl | pZ073 | ssDNA, Seq from Michaels et al.[2] | Yes |
| pLP192 | AmCyan-TRE-DsRed-V15 | pZ073 | ssDNA, Seq from Michaels et al.[2] | Yes |
| pLP193 | AmCyan-TRE-DsRed-T17 V9 | pZ073 | ssDNA, Seq from Michaels et al.[2] | Yes |

#### MoClo Level 0 backbones (to restore Kozak sequence)

| Name | Description | Backbone | Deposit on Addgene |
|---|---|---|---|
| pLP301 | 5' Kozak pICH41258 | pICH41258 | Yes |
| pLP302 | 5' Kozak pAGM1287 | pAGM1287 | Yes |
| pLP303 | 5'Kozak pICH41308 | pICH41308 | Yes |
| pLP220 | 3' Kozak pAGM1276 | pAGM1276 | Yes |
| pLP221 | 3' Kozak pICH41246 | pICH41246 | Yes |
| pLP222 | 3' Kozak pICH41295 | pICH41295 | Yes |

#### MoClo level 0 library

| Name | Description | Level 0 Backbone | Source of insert | Deposit on Addgene |
|---|---|---|---|---|
| pLP133 | UBC | pICH41233 | dsDNA, Seq from Schreiber et al, seq modified to make compatible for MoClo cloning | Yes |
| pLP134 | PIT2 | pICH41258 | dsDNA, Seq from Schreiber et al [3], codon modified for compatibility with MoClo cloning, originates from Weber et al. [4] | Yes |
| pLP135 | miR302a | pAGM1263 | PCR from pLP113, Seq from miRBase | Yes |
| pLP136 | miR302a | pAGM1299 | PCR from pLP113, Seq from miRBase | Yes |
| pLP137 | miR302b | pAGM1276 | PCR from pLP114, Seq from miRBase | Yes |
| pLP138 | miR302b | pAGM1301 | PCR from pLP114. Seq from miRBase | Yes |
| pLP139 | SV40 polyA | pICH41276 | PCR from pJS23, Schreiber et al. [3] | Yes |
| pLP140 | 5'UTR Spacer | pICH41246 | ssDNA | Yes |
| pLP142 | T375 4x | pAGM1299 | PCR from pLP112, Seq from miRBase | Yes |

| | | | | |
|---|---|---|---|---|
| pLP143 | T489 4x | pAGM1301 | PCR from pLP111, Seq from miRBase | Yes |
| pLP144 | T708 4x | pICH53388 | PCR from pLP110, Seq from miRBase | Yes |
| pLP145 | PolyA | pICH53399 | PCR from pLP103, polyA originates from pJS32, Schreiber et al. [3] | Yes |
| pLP146 | UBC | pICH41295 | PCR from pLP133, Seq originates from Schreiber et al. [3] | Yes |
| pLP147 | TFF6 4x | pAGM1299 | PCR from pLP119, Seq originates from Leisner et al.[1] | Yes |
| pLP148 | TFF6 4x | pAGM1301 | PCR from pLP119, Seq originates from Leisner et al.[1] | Yes |
| pLP149 | TFF6 4x | pICH53388 | PCR from pLP119, Seq originates from Leisner et al.[1] | Yes |
| pLP164 | TFF3 | pAGM1301 | PCR from pLP118, Seq originates from Leisner et al.[1] | Yes |
| pLP194 | TFF3 4x | pAGM1287 | ssDNA, Seq originates from Leisner et al.[1] | Yes |
| pLP195 | TFF6 4x | pAGM1287 | ssDNA, Seq originates from Leisner et al.[1] | Yes |
| pLP196 | T375 4x | pAGM1287 | ssDNA, Seq from miRBase | Yes |
| pLP197 | sBFP | pAGM1301 | PCR from pLP088/ pCS187, Stelzer et al.[5] | Yes |
| pLP198 | mCherry | pAGM1301 | PCR from pLP027/pKH026, Prochazka et al.[6] | Yes |
| pLP199 | iRFP | pAGM1301 | PCR from pLP029/pCS184, Prochazka et al.[6] | Yes |
| pLP206 | tTA | pICH41258 | PCR from Addgene plasmid #24415, codon modified for compatibility with MoClo cloning | Yes |
| pLP210 | ET1 | pICH41258 | dsDNA, seq from Prochazka et al [6], originates from Weber et al [7] | Yes |
| pLP223 | ETRtight | pAGM41295 | dsDNA, seq from Prochazka et al.[6], ETR binding sites originate from Weber et al. [7] | Yes |
| pLP233 | TFF4 3x | pICH41246 | dsDNA, Seq from Xie et al.[8] | Yes |
| pLP235 | TFF4 3x | pICH41264 | dsDNA, Seq from Xie et al[8] | Yes |
| pLP237 | tTAmut | pICH41258 | dsDNA, Seq from Roney et al.[9] | Yes |
| pLP238 | miR375 miR489 miR708 | pICH53388 | PCR from pLP169 | Yes |
| pLP240 | TRE3G | pICH41233 | PCR from pLP089, Seq originally from Addgene #61474 | Yes |
| pLP242 | 5’UTR Spacer (restores Kozak) | pLP221 | dsDNA | Yes |
| pLP243 | 5’UTR Spacer | pICH41246 | dsDNA | Yes |
| pLP245 | Citrine FF4 | pICH41258 with Kozak OH | PCR from pJS41, Schreiber et al [3] | Yes |
| pLP247 | 3’UTR spacer | pICH41264 | ssDNA | Yes |
| pLP248 | mCherry | pICH41258 | PCR from pLP027, Prochazka et al. [6] | Yes |
| pLP251 | PIRtight | pICH41233 | PCR from X, Prochazka et al.[6], PIR binding site originates from Weber et al. [4] | ? |
| pLP252 | ETRtight | pICH41233 | PCR from X Prochazka et al. [6], ETR binding site originally from Weber et al. [10] | ? |
| pLP253 | 5'UTR spacer | pICH41264 | ssDNA, Seq fom pLP027, Prochazka et al.[6] | Yes |
| pLP254 | rtTA | pICH41258 | PCR from pLP059 / Addgene #61472 | Yes |
| pLP255 | Ef1a | pICH41233 | PCR from pLP027, Prochazka et al.[6] | Yes |
| pLP257 | T375 4x | pAGM1299 | ssDNA, Seq from miRBase | Yes |
| pLP258 | T489 4x | pAGM1301 | ssDNA, Seq from miRBase | Yes |
| pLP259 | T708 4x | pICH53388 | ssDNA, Seq from miRBase | Yes |
| pLP305 | TFF3 4x | pAGM1299 | ssDNA, Seq from Leisner et al. [1] | Yes |
| pLP306 | TFF6 4x | pAGM1301 | ssDNA, Seq from Leisner et al. [1] | Yes |
| pLP314 | T302a 4x | pICH41264 | ssDNA, Seq from miRBase | Yes |
| pLP315 | TFF3 4x | pICH41264 | ssDNA, Seq from Leisner et al. [1] | Yes |

| | | | | |
|---|---|---|---|---|
| pLP316 | TFF4 3x | pAGM1299 | ssDNA, Seq from Xie et al. [8] | Yes |
| pLP317 | T375-3p | pAGM1301 | ssDNA, Seq from miRBase | Yes |
| pLP318 | T489 4x | pICH53388 | ssDNA, Seq from miRBase | Yes |
| pLP319 | TFF3 4x | pAGM1301 | ssDNA, Seq from Leisner et al. [1] | Yes |
| pLP320 | TFF6 4x | pICH53388 | ssDNA, Seq from Leisner et al. [1] | Yes |
| pLP332 | Spacer | pICH41264 | ssDNA | Yes |
| pLP333 | Spacer | pICH53388 | ssDNA | Yes |
| pLP334 | TRE3G | pLP222 | PCR from pLP089, Seq originates from Addgene #61474 | Yes |
| pLP337 | BMP4 | pLP301 | dsDNA, Seq from USCS, hg38_knownGene_ENST00000245451.9 range=chr14:53950032-53952222, CDS only, then codon optimized with IDT tool | Yes |
| pLP364 | PIRtight | pLP222 | PCR from pLP269, Seq from Prochazka et al.[6], binding site originates from Weber et al. [4] | Yes |
| pLP365 | mCherry | pLP301 | PCR from pLP027, Prochazka et al.[6] | Yes |
| pLP366 | sBFP2 | pLP301 | PCR from pLP027, Prochazka et al.[6] | Yes |
| pLP367 | T17 4x | pICH53388 | ssDNA, Seq from miRBase | Yes |
| pLP368 | T17 V10 | pICH53388 | ssDNA, Seq from Michaels et al. [2] | Yes |
| pLP349 | T17 V2 | pICH53388 | ssDNA, Seq from Michaels et al. [2] | Yes |
| pLP350 | T17 V4 | pICH53388 | ssDNA, Seq from Michaels et al. [2] | Yes |
| pLP351 | T17 V8 | pICH53388 | ssDNA, Seq from Michaels et al. [2] | Yes |
| pLP352 | T17 V1 | pICH53388 | ssDNA, Seq from Michaels et al. [2] | Yes |
| pLP353 | T17 V7 | pICH53388 | ssDNA, Seq from Michaels et al. [2] | Yes |
| pLP354 | T17 V14 | pICH53388 | ssDNA, Seq from Michaels et al. [2] | Yes |
| pLP355 | neg ctrl | pICH53388 | ssDNA, Seq from Michaels et al. [2] | Yes |
| pLP356 | T17 V12 | pICH53388 | ssDNA, Seq from Michaels et al. [2] | Yes |
| pLP357 | T17 V9 | pICH53388 | ssDNA, Seq from Michaels et al. [2] | Yes |
| pLP358 | T17 V5 | pICH53388 | ssDNA, Seq from Michaels et al. [2] | Yes |
| pLP359 | T17 V11 | pICH53388 | ssDNA, Seq from Michaels et al. [2] | Yes |

#### MoClo level1 library (functional plasmids)

| **Name** | **Description** | **Level 1 Backbone** | **Deposit to Addgene** |
|---|---|---|---|
| pLP264 | UBC-TFF4-ET1-TFF4-T375-T489-T708-polyA | pICH47802 | Yes |
| pLP265 | UBC-TFF4-PIT2-TFF4-T375-T489-T708-polyA | pICH47802 | Yes |
| pLP266 | UBC-TFF4-tTA-TFF4-T375-T489-T708-polyA | pICH47802 | Yes |
| pLP267 | UBC-TFF4-tTAmut-TFF4-T375-T489-T708-polyA | pICH47802 | Yes |
| pLP268 | ETRtight-mCherry-polyA | pICH47742 | Yes |
| pLP269 | PIRtight-mCherry-polyA | pICH47742 | Yes |
| pLP271 | TRE3G-mCherry-polyA | pICH47742 | Yes |
| pLP275 | TRE3G-Citrine FF4-polyA | pICH47742 | Yes |
| pLP292 | Ef1a-spacer-rtTA-TFF6-TFF3-polyA | pICH47802 | Yes |
| pLP293 | Ef1a-spacer-rtTA-T302a-T302b-polyA | pICH47802 | Yes |
| pLP294 | Ef1a-spacer-PIT2-TFF6-TFF3-polyA | pICH47802 | Yes |
| pLP295 | Ef1a-spacer-PIT2-T302a-T302b-polyA | pICH47802 | Yes |
| pLP299 | PIRtight-mCitrine FF4-polyA | pICH47742 | Yes |
| pLP324 | Ef1a-spacer-rtTA-T302a-polyA make map | pICH47802 | Yes |
| pLP325 | Ef1a-spacer-rtTA-FF3-polyA make map | pICH47802 | Yes |

| | | | |
|---|---|---|---|
| pLP327 | UBC-TFF4-PIT2-TFF4 T375 T489 make map | pICH47742 | Yes |
| pLP342 | TRE3G-iRFP-spacer-polyA | pICH47742 | Yes |
| pLP343 | TRE3G-BMP4-spacer-polyA | pICH47742 | Yes |
| pLP360 | UBC-TFF4-PIT2-TFF4 TFF3 TFF6 make map | pICH47742 | Yes |
| pLP370 | PIRtight_mCherry_spacer | pICH47802 | Yes |
| pLP371 | PIRtight_sBFP_spacer_polyA | pICH47822 | Yes |
| pCL039 | PIRtight-Cherry-Spacer-T17 WT 1x-polyA | pICH47802 | Yes |
| pCL040 | PIRtight-Cherry-Spacer-T17 V2-polyA | pICH47802 | Yes |
| pCL041 | PIRtight-Cherry-Spacer-T17 V4-polyA | pICH47802 | Yes |
| pCL042 | PIRtight-Cherry-Spacer-T17 V8-polyA | pICH47802 | Yes |
| pCL043 | PIRtight-Cherry-Spacer-T17 4x-polyA | pICH47802 | Yes |
| pCL044 | PIRtight-sBFP-Spacer-T17 WT 1x-polyA | pICH47822 | Yes |
| pCL045 | PIRtight-sBFP-Spacer-T17 V2-polyA | pICH47822 | Yes |
| pCL046 | PIRtight-sBFP-Spacer-T17 V4-polyA | pICH47822 | Yes |
| pCL047 | PIRtight-sBFP-Spacer-T17 V8-polyA | pICH47822 | Yes |
| pCL048 | PIRtight-sBFP-Spacer-T17 4x-polyA | pICH47822 | Yes |
| pCL056 | TRE3G-BMP4-Spacer-T17 WT 1x-polyA | pICH47742 | Yes |
| pCL057 | TRE3G-BMP4-Spacer-T17 V2-polyA | pICH47742 | Yes |
| pCL058 | TRE3G-BMP4-Spacer-T17 V4-polyA | pICH47742 | Yes |
| pCL059 | TRE3G-BMP4-Spacer-T17 V8-polyA | pICH47742 | Yes |
| pCL060 | TRE3G-BMP4-Spacer-T17 4x-polyA | pICH47742 | Yes |
| pCL112 | EF1a-spacer-tTA-T375 4x-T489 4x-polyA | pICH47802 | Yes |
| pCL113 | EF1a-spacer-tTA-T375 4x-T375 4x-polyA | pICH47802 | Yes |
| pCL114 | EF1a-spacer-tTA-T302a 4x-T302b 4x-polyA | pICH47802 | Yes |
| pCL115 | EF1a-spacer-tTA-TFF3 4x-TFF6 4x-polyA | pICH47802 | Yes |
| pCL116 | EF1a-spacer-PIT2-T375 4x-T489 4x-polyA | pICH47802 | Yes |

Other plasmids used in this study (not cloned)

| **Name** | **Description** | **Source** | **Deposit to Addgene** |
|---|---|---|---|
| pLP026 / pKH025 | Ef1a-mCitrine | Prochazka et al [6] | No |
| pLP027 / pKH026 | Ef1a-mCherry | Prochazka et al [6] | No |
| pLP029 / pCS184 | Ef1a-iRFP | Prochazka et al [6] | No |
| pLP088 / pCS187 | Ef1a-sBFP2 | Stelzer et al [5] | No |
| pLP076 / pJS37 | UBC-TFF4-mCherry-TFF4-polyA | Schreiber et al [3] | No |
| pLP085 / pZ145 | AmCyan-TRE-DsRed-T17 | Xie et al. [8] | No |
